# Spatial hepatocyte plasticity of gluconeogenesis during the metabolic transitions between fed, fasted and starvation states

**DOI:** 10.1101/2024.04.29.591168

**Authors:** Junichi Okada, Austin Landgraf, Alus M. Xiaoli, Li Liu, Maxwell Horton, Victor L. Schuster, Fajun Yang, Simone Sidoli, Yunping Qiu, Irwin J. Kurland, Carolina Eliscovich, Kosaku Shinoda, Jeffrey E. Pessin

## Abstract

The liver acts as a master regulator of metabolic homeostasis in part by performing gluconeogenesis. This process is dysregulated in type 2 diabetes, leading to elevated hepatic glucose output. The parenchymal cells of the liver (hepatocytes) are heterogeneous, existing on an axis between the portal triad and the central vein, and perform distinct functions depending on location in the lobule. Here, using single cell analysis of hepatocytes across the liver lobule, we demonstrate that gluconeogenic gene expression (*Pck1* and *G6pc*) is relatively low in the fed state and gradually increases first in the periportal hepatocytes during the initial fasting period. As the time of fasting progresses, pericentral hepatocyte gluconeogenic gene expression increases, and following entry into the starvation state, the pericentral hepatocytes show similar gluconeogenic gene expression to the periportal hepatocytes. Similarly, pyruvate-dependent gluconeogenic activity is approximately 10-fold higher in the periportal hepatocytes during the initial fasting state but only 1.5-fold higher in the starvation state. In parallel, starvation suppresses canonical beta-catenin signaling and modulates expression of pericentral and periportal glutamine synthetase and glutaminase, resulting in an enhanced pericentral glutamine-dependent gluconeogenesis. These findings demonstrate that hepatocyte gluconeogenic gene expression and gluconeogenic activity are highly spatially and temporally plastic across the liver lobule, underscoring the critical importance of using well-defined feeding and fasting conditions to define the basis of hepatic insulin resistance and glucose production.

## INTRODUCTION

The liver plays numerous roles in the control of normal physiology including the production of bile acids necessary for lipid absorption, secretion of NaHCO_3_ to neutralized gastric acid in the intestines, detoxification of endogenous and exogenous toxins, removal of excess ammonia by conversion to urea among other key physiologic process^1,2,3,4,5^. In particular, the liver generates glucose via gluconeogenesis and glycogenolysis during states of nutrient deprivation and conversely in states of nutrient excess repletes its glycogen stores, stimulates de novo lipogenesis for fatty acid synthesis from carbon precursors and packages newly synthesized lipids into very low-density lipoprotein particles for secretion and storage in adipose tissue^6,7,8^.

Dysregulation of these liver glucose and lipid metabolic processes are primary hallmarks of insulin resistance and dyslipidemia that occur in obesity and type 2 diabetes^9,10^. Although the parenchymal cells of the liver (hepatocytes) comprise approximately 50% of the liver cells (∼70% based on mass), they are uniquely responsible for the activation of glucose production in the fasted state and for activation of de novo lipogenesis in the fed state^11,12,13,14^. Many studies have further shown that hepatocytes have distinct functions across the periportal to pericentral spatial axis within each individual lobule^15,16,17^. For example, glycogen synthesis^18^ and gluconeogenesis^19,20,21^ have both been shown to be biased towards the periportal zone. Although two studies have reported that de novo lipogenesis is pericentrally biased^22,23^, and one study has suggested that this process may be more prominent in periportal hepatocytes^24^.

In any case, the normal physiologic temporal regulation of gluconeogenesis and de novo lipogenesis across the liver lobule during the transitions from the fed to fasted state and subsequent fasted to starvation state are important and unanswered questions, as progress in understanding the bases for metabolic pathophysiology is not possible without first understanding normal physiologic regulation. Here, with the advent of single cell and spatial technologies, and zonation specific hepatocyte cell isolation, we have undertaken a systematic approach to analyze the in vivo temporal and spatial patterns of gluconeogenic gene expression in correlation with functional analysis of gluconeogenesis activity across the liver lobule during the feeding/fasting/starvation metabolic transitions. These data demonstrate a remarkable hepatocyte plasticity of gluconeogenic gene expression and gluconeogenic activity across the liver lobule depending upon organismal nutritional state. Our results suggest this hepatic plasticity results from a fasting induced suppression of canonical β-catenin (WNT signaling) resulting in a decline of hepatocyte zonation with the pericentral hepatocytes taking on periportal gene expression characteristics.

## RESULTS

### Time restricted feeding regulation of gluconeogenic and lipogenic gene expression

To investigate the regulation of liver gene expression in the normal physiologic transitions between the fed, fasted and starvation states, we established a paradigm in which mice rapidly consume food and saturate their stomachs to create a relatively uniform fed state. Removal of food after 4h of feeding facilitates a relatively uniform transition to the fasted and then starvation states. This was accomplished by allowing mice to access standard low-fat laboratory chow (20 kcal% protein, 4.5 kcal% fat) between 9:00 and 17:00 each day for 3 days. After these 3 days of training on the 4th day at 9:00, the mice were provided food and 20% sucrose water for 4h as previously described^25,26^. The 0h (fully fed) time point was defined as 13:00 (Fig. 1A). At 13:00 the food was removed, and the sucrose water was replaced with regular water. Mice were subsequently fasted for indicated time periods (21:00 = 8h, 5:00 = 16h, 13:00 = 24h, 19:00 = 30h following food removal. To confirm the effectiveness of this protocol, the relative induction of *Fasn* and *Pck1* mRNA was used to assess metabolic state (Fig. S1A). At the start of feeding (−4h), the hepatocytes display a relatively high expression level of the gluconeogenic gene *Pck1* and low levels of the lipogenic gene *Fasn* (Fig. S1B and S1C). *Pck1* mRNA expression starts to decline at the −2h time point, and at approximately the same time, *Fasn* mRNA starts to increase. At the 0h time point, *Pck1* mRNA is substantially reduced while *Fasn* mRNA is substantially increased. Following the removal of food and sucrose water (0h), lipogenic gene expression (*Fasn*) is substantially reduced by 4h and further declines at 24h (Fig. S1D). In contrast, gluconeogenic gene expression (*Pck1*) starts to increase at 4h and further increases at 24h (Fig. S1E). In parallel, we analyzed changes in liver glycogen content (Fig. S1F).

**Fig. 1.**
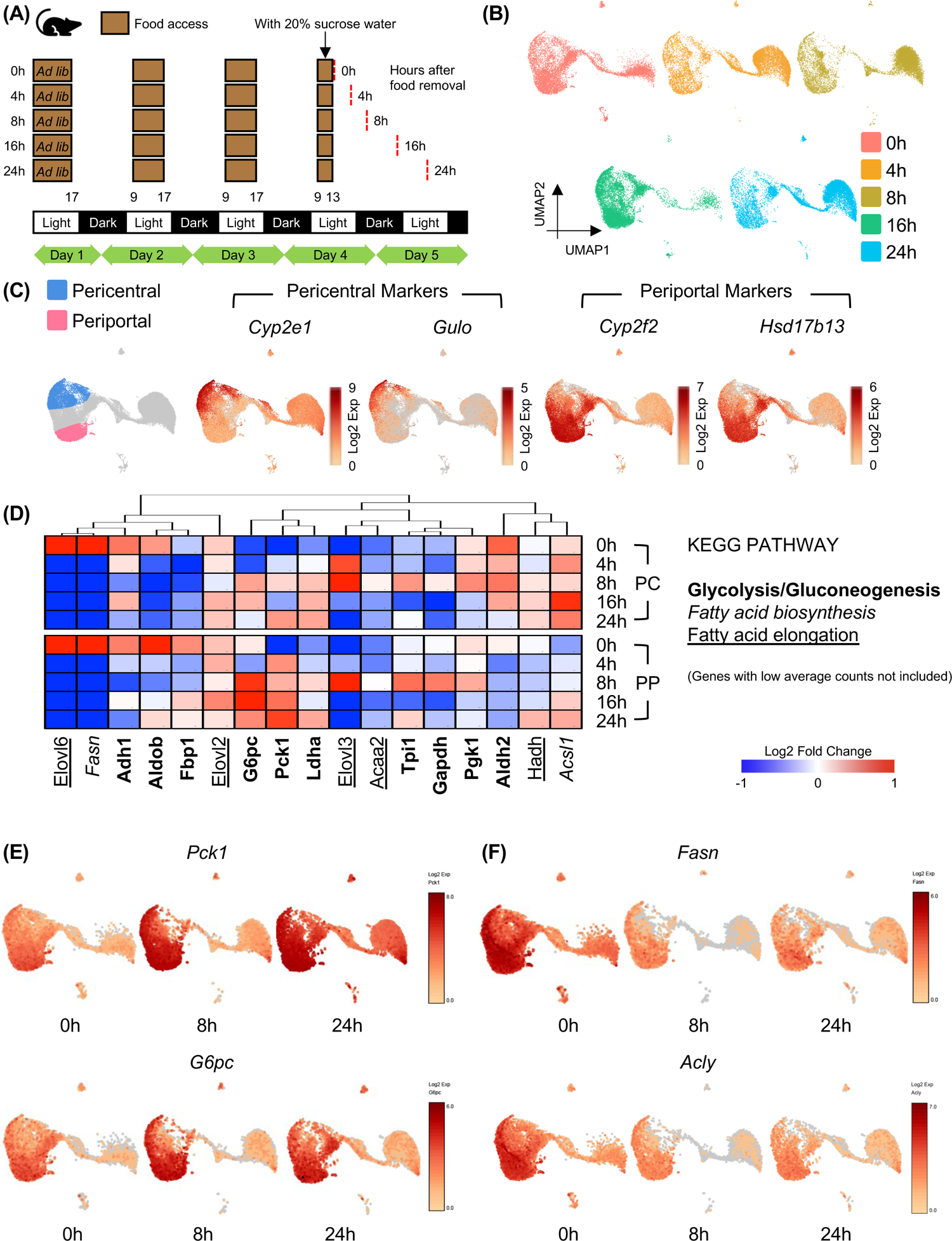
Standard scRNA-seq reveals zonation and activated pathways during fed/fasting. **(A)** Scheme of the feeding and fasting protocol. **(B)** UMAP visualizations of the standard scRNA-seq based on feeding conditions; fed4h (0h), fast4h (4h), fast8h (8h), fast16h (16h), and fast24h (24h) (N=1 mouse per time point, n=50,000 hepatocytes total, each dots represents one cell). **(C)** Pericentral and periportal hepatocyte identification based on the expression levels of pericentral zonation markers (*Cyp2e1* and *Gulo*) and periportal zonation markers (*Cyp2f2* and *Hsd17b13*). **(D)** Heatmap of metabolic related KEGG PATHWAYs. Glycolysis/Gluconeogenesis genes are in bold, fatty acid biosynthesis genes are in italic, fatty acid elongation genes are underlined. Pericentral and periportal hepatocytes are clustered based on Fig. 1C. Genes with low average counts are not included. **(E, F)** UMAP plots of gluconeogenic genes (*Pck1*, *G6pc*) and Lipogenic genes (*Fasn*, *Acly*) in the fed (0h), fasted (8h), and starvation (24h) state.

There was no reduction in glycogen content for up to 8h of food deprivation, but at 16h glycogen was reduced approximately 60%, and by 24h it was essentially zero. Thus, we consider food deprivation for up to 16h as a fasted state and food deprivation beyond 16h as a starvation state. This data demonstrates that our feeding/fasting protocol results in a uniform and controlled transition between the fed, fasted and starvation states for individual mice.

### Spatial and temporal regulation of liver lobule hepatocyte gene expression

Having established a controlled model of nutritional states, we utilized the 10x Genomics single cell RNA-seq (scRNA-seq) platform in primary isolated hepatocytes to examine the expression profiles of periportal and pericentral hepatocytes. Seurat was used to establish identical UMAP clusters of hepatocytes in the fully fed (0h), fasted (4 to 8h) and starvation (16 to 24h) states (Fig. 1B). To identify the position of the periportal and pericentral hepatocytes, we took advantage of previous studies that have characterized the localization of specific zonation markers^27,28^. The pericentral markers *Cyp2e1* and *Gulo* were highly colocalized to one region while the periportal markers *Cyp2f2* and *Hsd17b13* were highly colocalized to a different region (Fig. 1C). As these genes are typically expressed as a gradient across the lobule, the cells that have low expression of both pericentral and periportal markers likely represent the mid-zone hepatocytes. As expected, gluconeogenic genes, such as *Pck1,* are generally restricted to the periportal hepatocytes, induced during the fed/fasted transition, and maintained in the starvation states (Fig. 1D). However, transition into the starvation state also results in the induction of gluconeogenic gene expression in the pericentral hepatocytes. In contrast, lipogenic genes, such as *Fasn*, are expressed in both the pericentral and periportal hepatocytes in the fed state, but after food removal, these genes are rapidly shut off and remain off over the entire fast/starvation course (Fig. 1D). The UMAP plots for the time course of *Pck1* and *G6pc* mRNAs are shown in Figure 1E, and the plots for *Fasn* and *Acly* mRNAs are shown in Figure 1F.

COMPASS is an algorithm that derives information on cellular metabolic states from single-cell transcriptomic data^29^. Using scRNA-seq data combined with a flux balance model based on RECON2, COMPASS can generate predictions on flux dynamics to determine the differential potential-activity of various metabolic pathways in the periportal and pericentral hepatocytes. Compared to the fed state, 8h of fasting resulted in a marked activation of periportal metabolic pathways (Fig. 2A) that was somewhat further increased following 24h of fasting (Fig. 2B). The top 5 activated pathways at 8h (highlighted in Fig. 2) remained highly activated at 24h in periportal hepatocytes. In contrast, there was no significant activation of pericentral hepatocyte metabolic pathways between the fed state and following 8h of fasting (Fig. 2C). However, by 24h of fasting the pericentral hepatocytes displayed a strong activation of metabolic pathways similar to that of the periportal hepatocytes (Fig. 2D). Together, these data indicate that periportal hepatocytes become metabolically more active upon entry into the fasted state whereas pericentral hepatocyte remain relatively refractory. However, upon entry into the starvation state the pericentral hepatocytes become highly metabolically active supporting a temporal conversion of pericentral hepatocytes to a more periportal phenotype.

**Fig. 2.**
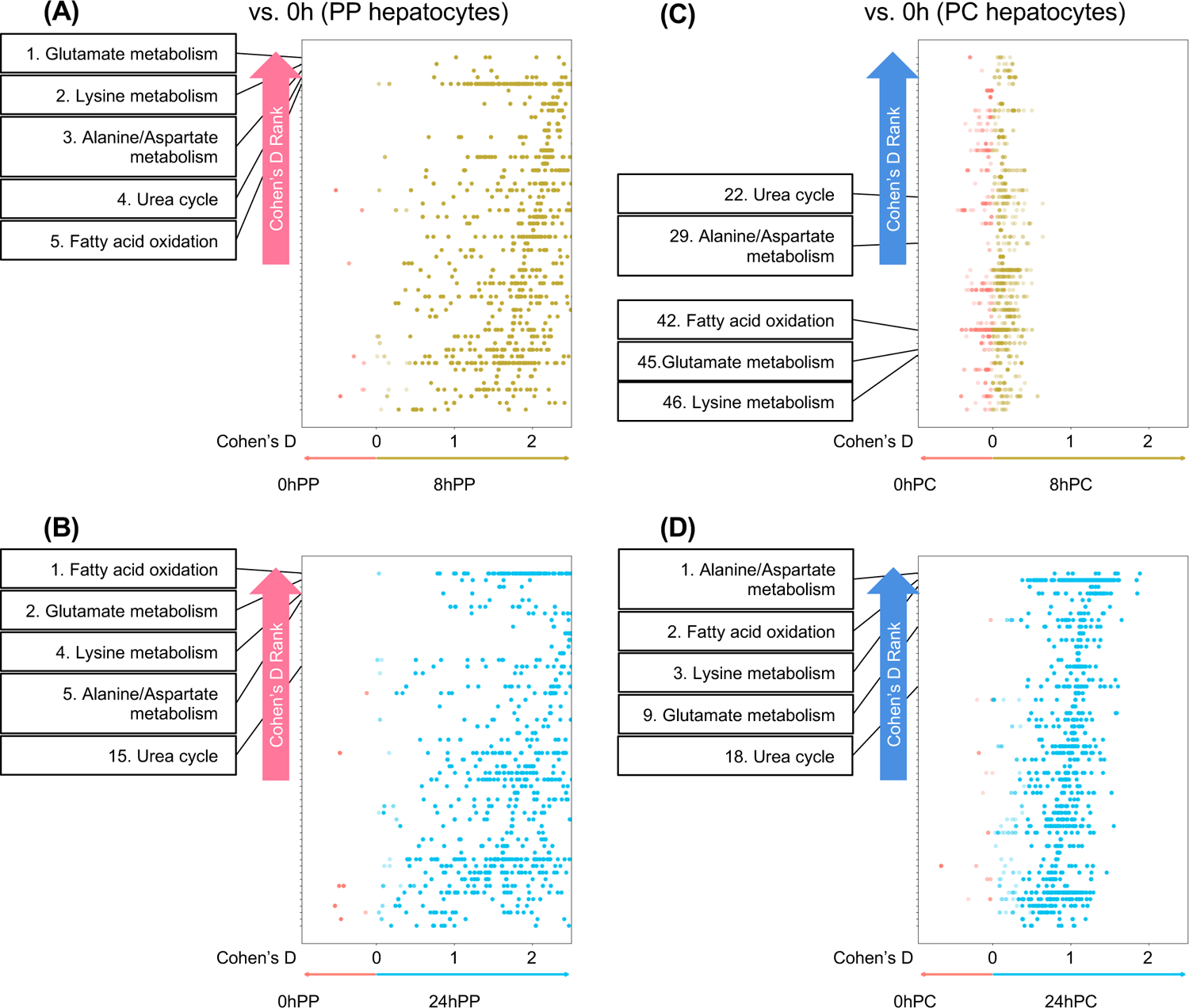
Metabolic flux increases in pericentral hepatocytes in the starvation state. **(A-D)** Compass^29^ analysis comparison between pericentral and periportal fed (0h=red), fasted (8h=yellow), and starvation (24h=blue) state on standard scRNA-seq. Each reaction (dot) is colored/plotted based on Cohen’s D statistic and by Recon2 pathways. Top 5 activated pathway in 8h periportal hepatocytes are highlighted. Numbers show how high each pathway is upregulated by fasting (rankings based on Cohen’s D). **(A)** Comparison of differential activity of metabolic reactions between 0h periportal and 8h periportal hepatocytes. **(B)** Comparison of differential activity of metabolic reactions between 0h periportal and 24h periportal hepatocytes. **(C)** Comparison of differential activity of metabolic reactions between 0h pericentral and 8h pericentral hepatocytes. **(D)** Comparison of differential activity of metabolic reactions between 0h pericentral and 24h pericentral hepatocytes.

### Quantitative targeted single cell gluconeogenic and lipogenic gene expression

Although standard scRNA-seq data indicate plasticity of hepatocytes across the periportal to pericentral lobule, this technology is not designed for quantitative comparisons of individual gene expression patterns, and in particular for genes with medium-low expression levels^30^. To address this issue, we first performed quantitative single cell targeted RNA-seq for 61 specific genes consisting of pericentral and periportal hepatocyte markers, general parenchymal markers, non-parenchymal cell markers, gluconeogenic genes, lipogenic genes, bile acid synthesis genes, and several secreted hepatokines (Table 1). Saturation kinetics (the metric quantifies the fraction of reads that originate from a previously observed UMI) clearly demonstrated that standard scRNA-seq was unable to reach a 90% sequence saturation threshold and only was able to capture ∼70% of transcripts at each time point (Fig. S2A). In contrast, targeted scRNA-seq was able to capture ∼95% of the total number of unique transcripts present in each library. Comparison of the fed (0h) and fasted (16h) time points between the standard and targeted scRNA-seq datasets demonstrates a more robust regulation of *Pck1* and *Fasn* mRNA expression in the targeted single cell analysis (Fig. S2B).

**Table 1.**
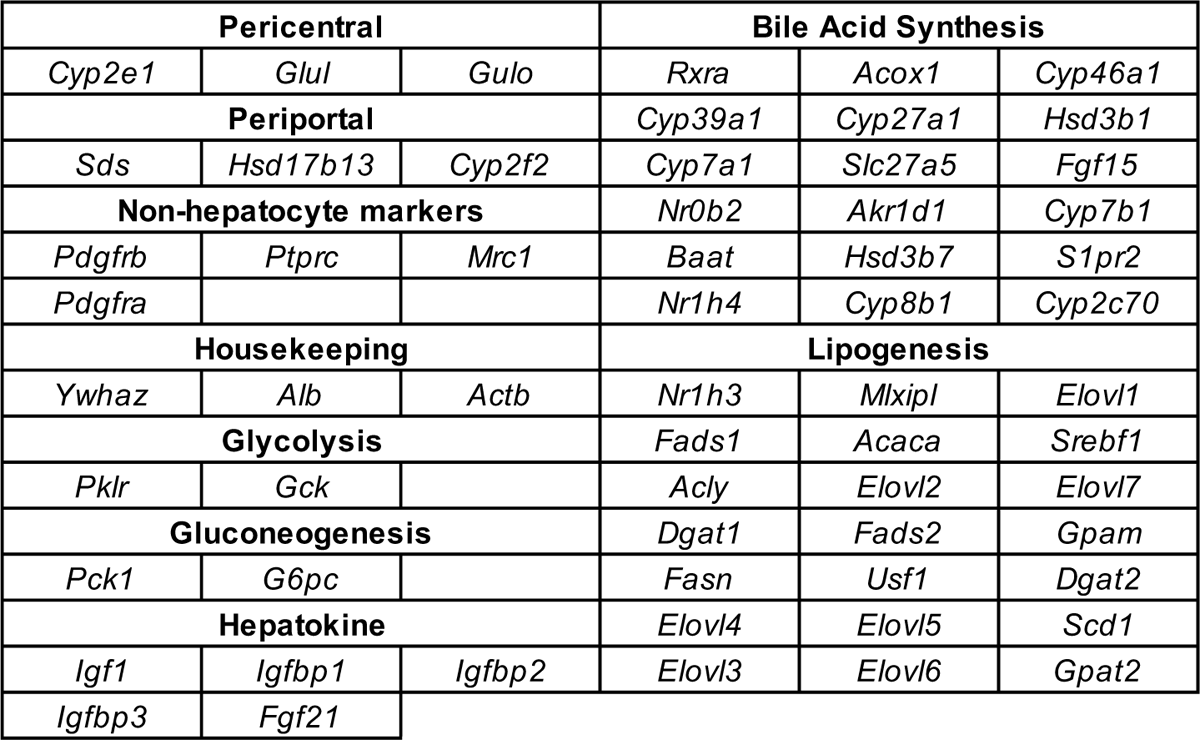
Gene list for the targeted gene expression analysis.

Combining all time points into a single two-dimensional UMAP plot for the entire feeding/fasting time course clearly demonstrates that the fed state (0h) is the most divergent cluster. In contrast, the fasted/starvation time points closely overlap each other with the 8h fasted and 30h starved state being the most divergent among the food restricted time points (Fig. 3A). The identification of periportal and pericentral hepatocytes from the targeted scRNA-seq is shown in Figure 3B based upon the distribution of several periportal and pericentral markers, four of which are shown in Figure 3C. Quantification of the time dependent changes in the two major gluconeogenic genes in the targeted scRNA-seq indicate that as hepatocytes transition from the early fasted to late fasted/starvation states, the periportal hepatocytes rapidly express *Pck1* and *G6pc* mRNA (Fig. 3D). Nevertheless, the pericentral hepatocytes also increase their expression of these genes, and in the case of *Pck1*, expression is nearly equal in pericentral and periportal hepatocytes after 30h of food restriction (Fig. 3D). In contrast, the majority of lipogenic genes are approximately equally expressed in periportal and pericentral hepatocytes in the fed (0h) state, and rapidly decline in parallel following food removal, becoming nearly undetectable by 8h of fasting (Fig. 3E). This divergence of expression patterns between the gluconeogenic and lipogenic genes is more apparent when displayed as ridge plots (Fig. S2C). Using the methods of Olsen et al.^31^, we performed fuzzy c-means (FCM) clustering to analyze the expression patterns of other genes. The target genes were clustered into 8 separate categories; those that are early fasting induced, late fasting/starvation induced, early fasting suppressed, late fasting/starvation suppressed, constitutive and cycling (2 other categories (other and low expressed) are not shown (Fig. S3).

**Fig. 3.**
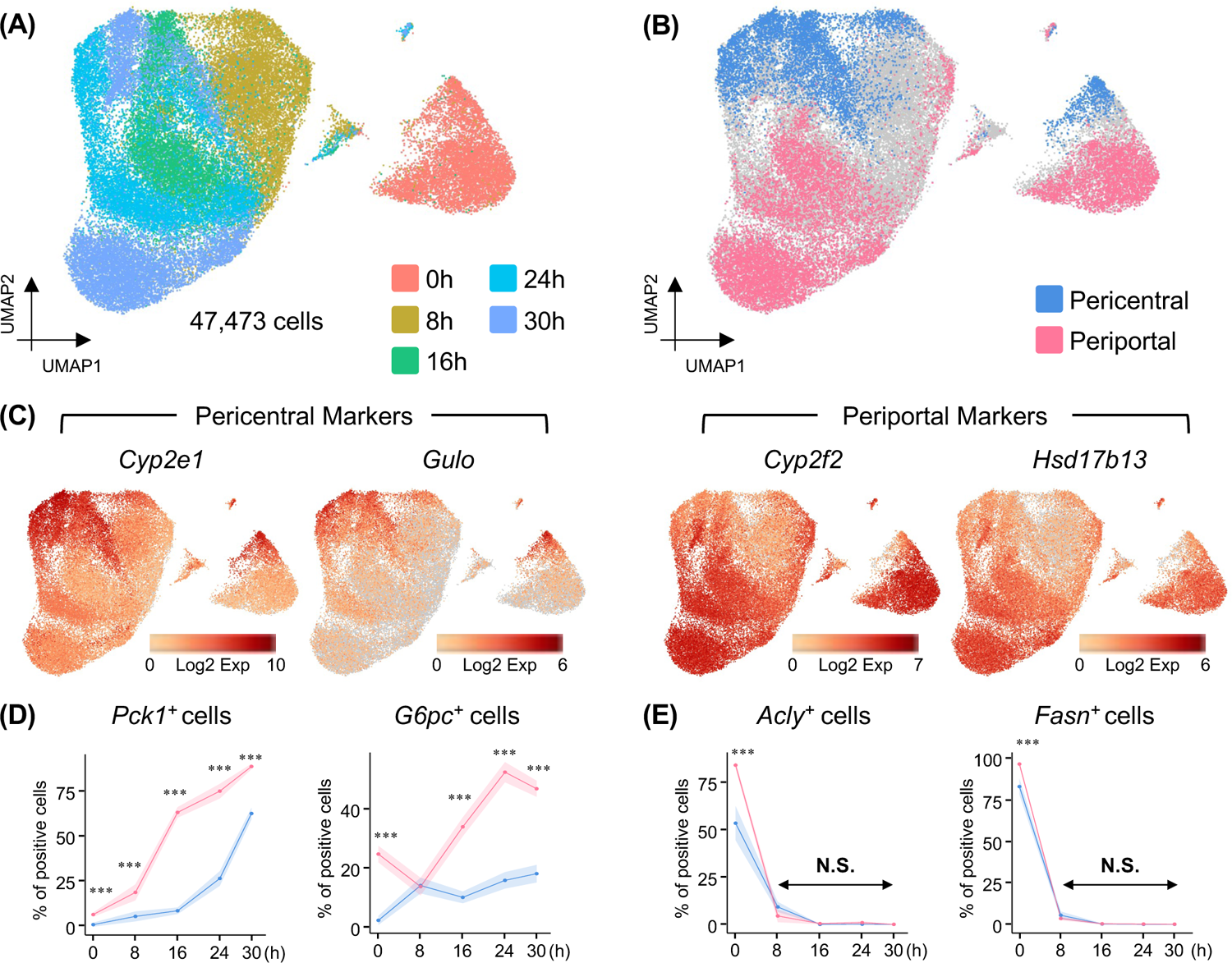
Unique kinetics of the gluconeogenic gene activation in single hepatocytes. **(A)** UMAP visualizations of the targeted scRNA-seq based on feeding conditions; fed4h (0h), fast8h (8h), fast16h (16h), fast24h (24h), and fast30h (30h) (N=1 mouse per time point, n=47,473 hepatocytes total, each dots represents one cell). **(B, C)** Pericentral and periportal hepatocyte identification based on the expression levels of pericentral zonation markers (*Cyp2e1* and *Gulo*) and periportal zonation markers (*Cyp2f2* and *Hsd17b13*). **(D, E)** Quantitative analysis based on the percentage of gluconeogenic (*Pck1*, *G6pc*) and lipogenic (*Fasn*, *Acly*) gene positive cells in the pericentral and periportal hepatocytes throughout the feeding and fasting time course. X-axis represents time points; Y-axis represents percentage of positive cells.

### Quantification of time restricted feeding on hepatocyte gluconeogenic and lipogenic gene transcription

We confirmed these findings using single-molecule Fluorescence *in situ* Hybridization (smFISH), which enables a direct visualization of individual mRNA molecules within the cell in the context of the liver tissue while preserving morphology. In the fed state (0h), a small fraction of periportal, and to a lesser extent pericentral hepatocytes that were dimmer for *Pck1* mRNA (Fig. 4A). Following 4h of fasting, the periportal and mid-zone hepatocytes displayed a marked increase in mRNA, with a smaller increase in the pericentral hepatocytes. As the time of fasting increased and transitioned into the starvation state (24h), the expression of *Pck1* increased in both the periportal and pericentral hepatocytes. Single-molecule FISH allowed the visualization of active nuclear transcription sites (TS) of the *Pck1* gene (Fig. 4A). We quantified the number of TS per liver zone using FISH-quant by selecting images centered around either the central vein or the portal triad (see Methods)^32,33^. Representative images of images used for quantification are shown in Figures S4A (for *Pck1*) and S4B (for *Fasn*). In most cases, nuclei display two discrete transcription sites that correspond to the two *Pck1* alleles. However, since hepatocytes are polyploid, some nuclei display more than two transcribing alleles. In addition, transcription occurs in bursts^34,35^, and therefore there are some nuclei with no detectable transcription sites, suggesting that *Pck1* transcription was dormant at this particular point in time (Fig. 4A). Consistent with the targeted scRNA-seq, quantification of the number of active TS per nuclei showed an increase first in the periportal hepatocytes and subsequently in the pericentral hepatocytes during the feeding/fasting transition (Fig. 4B). However, there was no difference in the intensity of individual transcription sites (Fig. S5A).

**Fig. 4.**
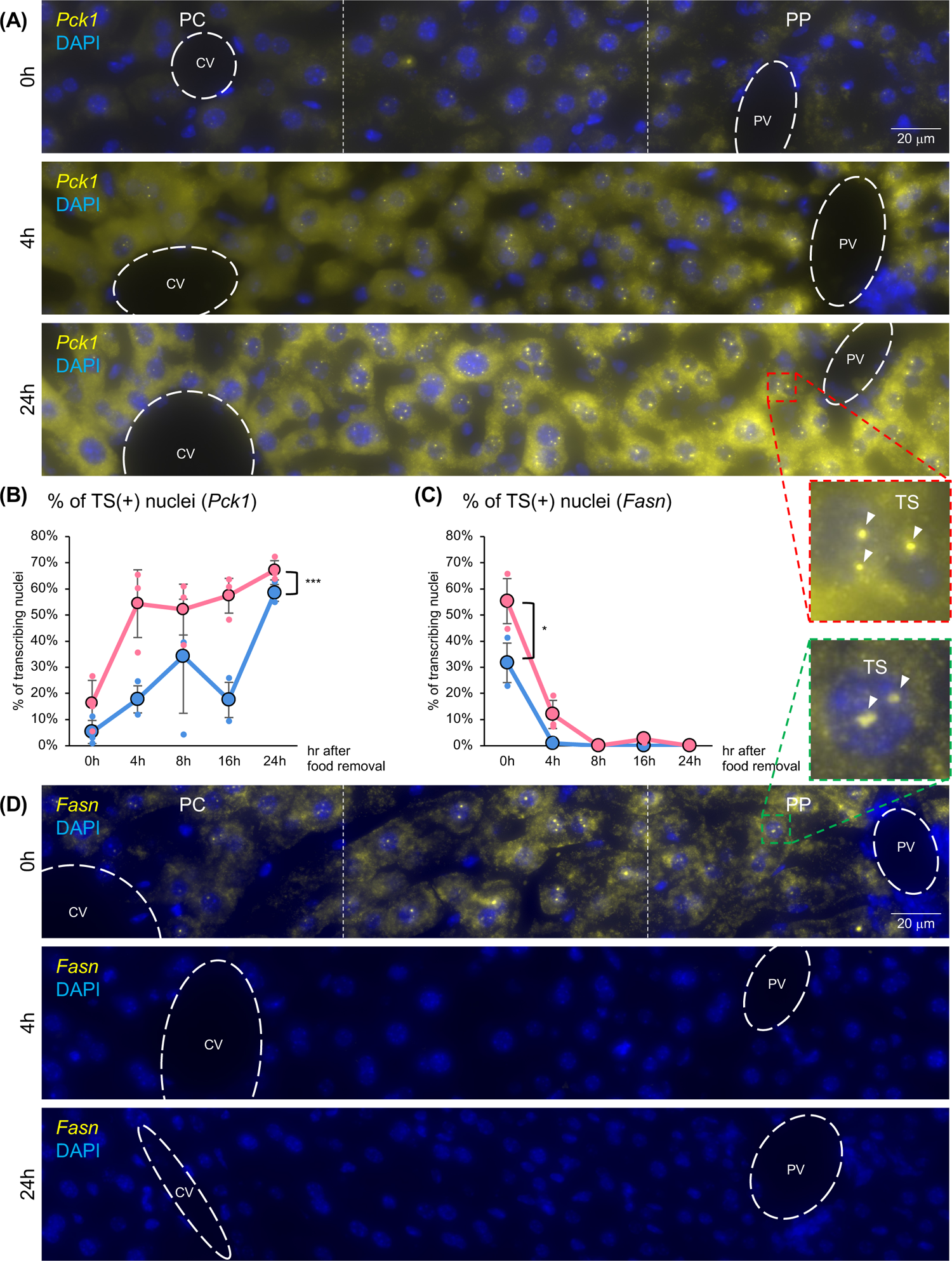
Transcription dynamics of gluconeogenic and lipogenic genes are different. **(A, D)** smFISH image of *Pck1* **(A)** and *Fasn* **(D)** in the fed (0h), fasted (4h), and starvation (24h) state (representative image shown for each time point, N=3 mice per time point). Lobules always show the central vein (CV) on the left, and the portal vein (PV) on the right. Magnified images show nuclei that have transcription sites (TS). **(B, C)** Quantitative analysis of TS by FISH-QUANT (minimum of 800 nuclei in 6 veins were analyzed per time point and zone) based on the percentage of *Pck1* **(B)** and *Fasn* **(C)** transcribing nuclei in the pericentral and periportal hepatocytes throughout the feeding and fasting time course. X-axis represents time points; Y-axis represents percentage of transcribing nuclei (dots highlighted with black rim show average of N=3, while other dots represent each mouse). Statistics based on repeated measures ANOVA.

In contrast, *Fasn* mRNA FISH signal was distributed across the lobule in the fed state with both periportal and pericentral hepatocytes displaying TS in their nuclei. *Fasn* signal was approximately 1.8-fold higher in the periportal hepatocytes (Fig. 4C and D).

Following 4h of fasting, there were barely any detectable *Fasn* mRNA or active TS by smFISH; by 24h of food deprivation, the smFISH signal was completely absent (Fig. 4D). The number of active *Fasn* TS per nuclei showed a rapid decline following the initiation of the fasted state in both pericentral and periportal hepatocytes (Fig. 4C and S5B). These findings suggest that the transcriptional dynamics of gluconeogenic and lipogenic genes are distinct, and that gluconeogenesis exhibits both temporal and spatial aspects of heterogeneity.

### Gluconeogenic activity in isolated primary periportal and pericentral hepatocytes

To assess the functional consequences of the observed temporal and spatial regulation of gluconeogenic gene expression in periportal and pericentral hepatocytes, we first utilized the digitonin ablation methodology^19,36,37^ to isolate periportal and pericentral hepatocytes, enabling comparison to total primary isolated hepatocytes. The relative purity and separation of the periportal and pericentral hepatocytes was confirmed using RT-qPCR and exhibited an average 17-fold enrichment of *Cyp2e1* mRNA in the isolated pericentral hepatocytes and a corresponding 19-fold enrichment of *Cyp2f2* mRNA in the isolated periportal hepatocytes (Fig. 5A). RT-qPCR of *Pck1* and *G6pc* mRNA from the isolated periportal and pericentral hepatocytes was consistent with the single cell (standard and targeted) RNA-seq and smFISH findings, supporting a greater increase of gluconeogenic gene expression in the periportal hepatocytes first followed by the pericentral hepatocytes (Fig. 5B). Deep RNA-seq at the 0, 8 and 24h time points from isolated periportal and pericentral hepatocytes also demonstrated consistent temporal and spatial trends for gluconeogenic and lipogenic genes (Fig. 5C). Immunoblotting for the *Pck1* gene protein product (phosphoenolpyruvate carboxykinase, PEPCK) showed higher protein levels in the fed periportal hepatocytes compared to pericentral hepatocytes (Fig. 5D and E). Moreover, PEPCK protein displayed a greater increase during the fasting time course in the periportal hepatocytes, but in the starvation state (24h), there was a substantial increase in PEPCK in the pericentral hepatocytes, approaching the protein levels in the periportal hepatocytes. Immunoblotting for Cyp2e1 (pericentral), E-cadherin (periportal), and vinculin (loading control) was used to confirm zonation at the protein level.

**Fig. 5.**
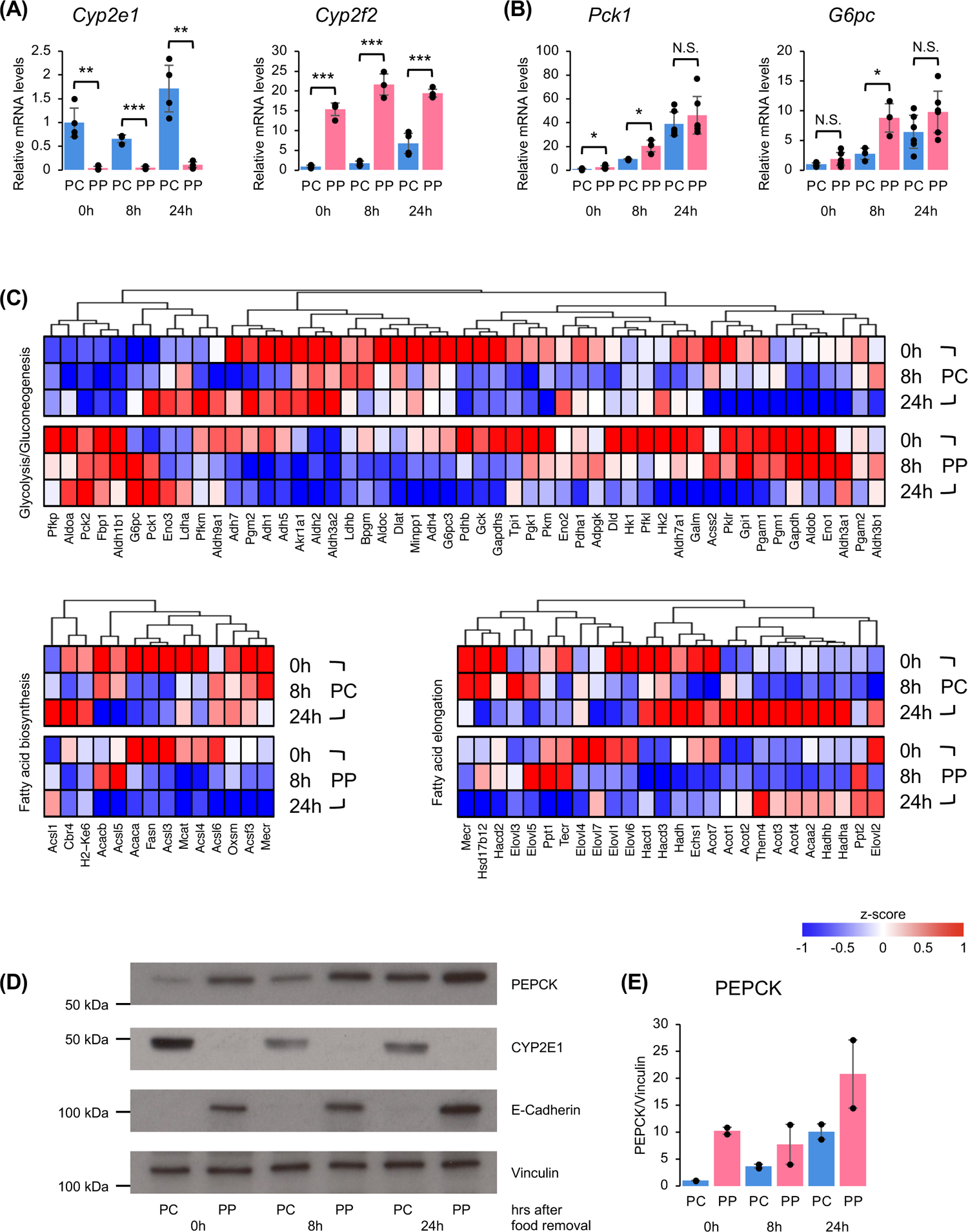
Digitonin ablation enables spatial and temporal heterogeneity analysis. **(A, B)** RT-qPCR (relative mRNA levels) of zonation markers (*Cyp2e1* pericentral, *Cyp2f2* periportal) **(A)** and gluconeogenic genes (*Pck1*, *G6pc*) **(B)** from pericentral and periportal isolated hepatocytes. N=3-6 mice per time point/condition. **(C)** Heatmap of metabolic related KEGG PATHWAYs (Glycolysis/Gluconeogenesis genes, fatty acid biosynthesis genes, fatty acid elongation genes) in RNA-seq from pericentral and periportal isolated hepatocytes. Average expression of N=3 from each time point/condition is shown. Coloring based on z-score. Genes with low average counts are not included. **(D)** Representative immunoblotting of PEPCK, CYP2E1, E-Cadherin, and Vinculin from pericentral and periportal isolated hepatocytes. **(E)** Quantification of PEPCK by Vinculin (N=2 mice per time point/condition).

We then assessed the functional gluconeogenic activity in isolated periportal and pericentral hepatocytes using stable isotope flux methodologies (Fig. 6A). We examined the time course of food deprivation on the formation and secretion of m2-glucose from 2,3[^13^C_2_]-pyruvate. For total isolated hepatocytes, there was no difference in media m2-glucose levels in the fed (0h) and fasted (8 and 16h) states (Fig. 6B). However, in the starvation state when glycogen is depleted (Fig. S1D), pyruvate-dependent glucose production was substantially increased. For isolated periportal and pericentral hepatocytes, both populations displayed relatively low m2-glucose production at 0h, 8h, and 16h, despite some variability amongst the pericentral samples at these time points (Fig. 6C). However, both the pericentral and periportal hepatocytes demonstrated a significant increase in m2-glucose production in the starved state (20 and 24h) with the periportal hepatocyte glucose production being approximately 1.5 to 2-fold greater than the pericentral hepatocyte production. However, when comparing the 0 and 24h time points for the pericentral hepatocytes, there is an approximate 10-fold increase in m2 glucose production, whereas the periportal hepatocytes only display an approximate 4-fold increase. These findings demonstrate that functional gluconeogenic activity can significantly increase in pericentral hepatocytes after prolonged food deprivation but remains higher in the periportal hepatocytes compared to the pericentral hepatocytes. A key decision point for whether substrate flux goes towards de novo lipogenesis or gluconeogenesis is the conversion of pyruvate to either acetyl CoA by pyruvate dehydrogenase (PDH) or to oxaloacetate by pyruvate decarboxylase (PC) ^38,39,40,41^.

**Fig. 6.**
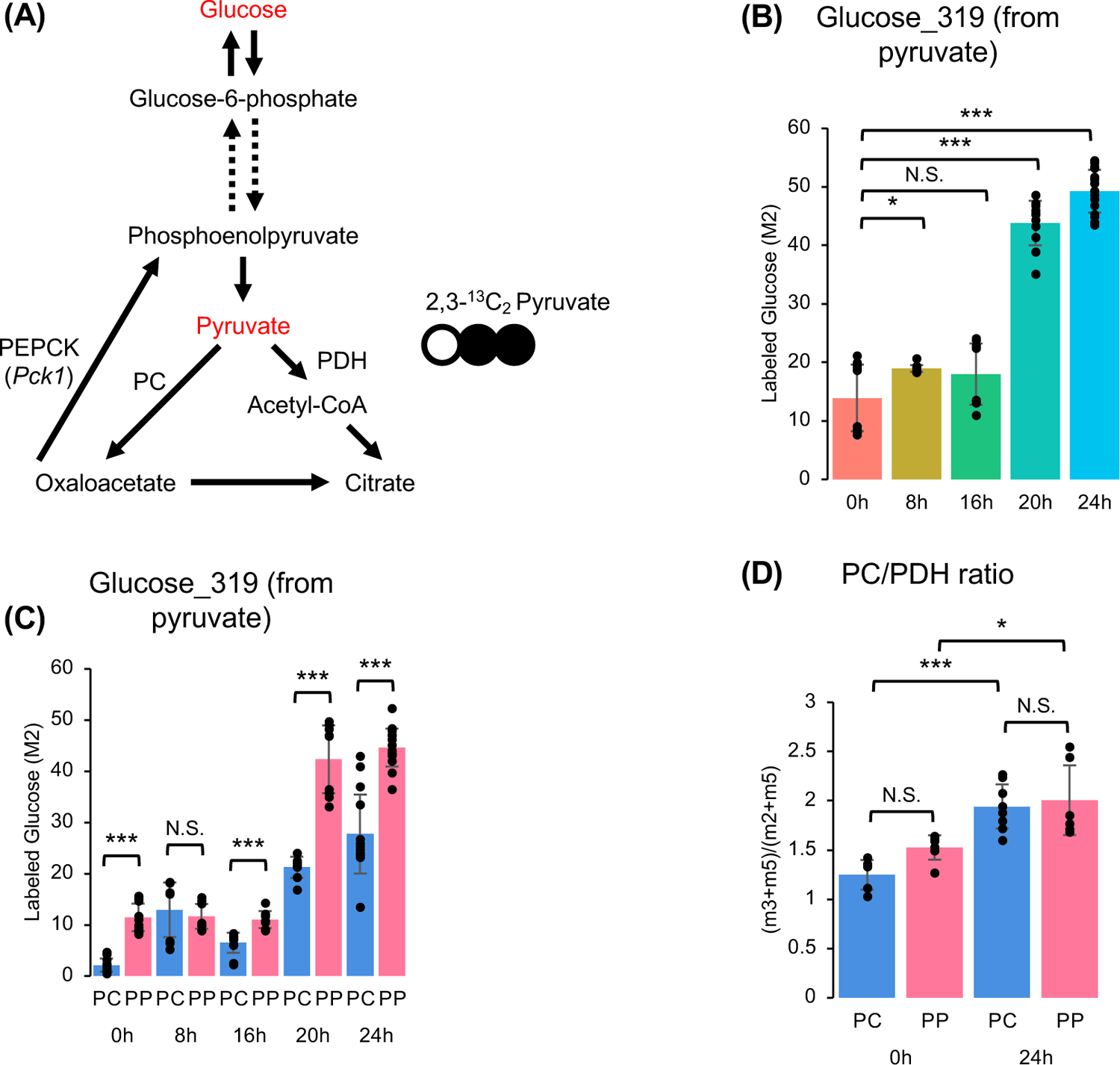
Gene expression directly regulates glucose production from pyruvate. **(A)** Scheme of 2,3 [^13^C_2_] pyruvate flowing into either pyruvate carboxylase (PC) or pyruvate dehydrogenase (PDH), and eventually to glucose. **(B)** Glucose_319 measurement from 2,3 [^13^C_2_] pyruvate within each time point from total hepatocytes. X-axis represents time points; Y-axis represents labeled glucose (M2) (dots represent N=2-4 mice per time point/condition combined with technical replicates). ANOVA followed by Tukey’s multiple comparisons test was performed to compare 0h with different time points. **(C)** Glucose_319 measurement from 2,3 [^13^C_2_] pyruvate within each time point from pericentral and periportal isolated hepatocytes. X-axis represents time points; Y-axis represents labeled glucose (M2) (dots represent N=2-5 mice per time point/condition combined with technical replicates). **(D)** PC/PDH ratio ((m3+m5)/(m2+m5) ratio of Citrate_465 from [^13^C_3_] pyruvate) within each time point from pericentral and periportal isolated hepatocytes. X-axis represents time points; Y-axis represents PC/PDH ratio (dots represent N=2 mice per time point/condition combined with technical replicates). ANOVA followed by Tukey’s multiple comparisons test was performed to compare 0h with different time points.

Examination of [^13^C_3_]-pyruvate flux in both pericentral and periportal isolated hepatocytes showed the expected increase in pyruvate carboxylase activity relative to pyruvate dehydrogenase activity following 24h of food deprivation (Fig. 6D).

### Fasting/starvation reduces pericentral hepatocyte canonical beta-catenin transcriptional activity

Hepatic zonation is primarily established by several signaling gradients across the liver lobule, with multiple selective WNT signaling target genes being highly expressed in the pericentral zone and progressively declining towards the periportal zone^42,43,44,45,46^. WNT signaling also negatively regulates some genes in the pericentral zone^42,45^. Anti-parallel to this, HIPPO/YAP and Sonic Hedgehog signaling for several selective target genes was reported to be generally higher in the periportal zone with progressive declines across the lobule towards the pericentral zone^47,48,49^. We analyzed RNA-seq data from periportal and pericentral hepatocytes at the 0h, 8h, and 24h time points to evaluate changes in WNT and HIPPO/YAP signaling during fasting. Consistent with RT-qPCR results, RNA-seq displayed the expected expression of zonation marker genes, gluconeogenic genes, and lipogenic genes during the fed/fasted/starvation transitions (Fig. S6A and B). Genome wide inspection of 103 WNT/β-catenin target genes indicated that 68% of WNT/β-catenin target genes temporally decline in the pericentral hepatocytes during the progression from the fed (0h) to fasted (8h) to starvation (24h) time points (Fig. S6C). Although a subset of annotated WNT/β-catenin target genes, such as *Gls2* and *Cdh1* (18 total), have relatively high expression in the periportal hepatocytes, these markedly declined in the fasted/starvation state compared to the fed state. Of the 103 target genes, there were a subset of genes that had increased expression in pericentral hepatocytes as well. The net changes in these canonical β-catenin target genes suggests that the pericentral hepatocytes are acquiring a periportal-like signature. Several canonical representative WNT/β-catenin target genes are presented in a more readily visualized quantitative manner in Figure S4D, as well as WNT (*Wnt5a* and *Wnt5b*) and β-catenin (*Ctnnb1*).

Examination of 39 HIPPO/YAP target genes indicated that several are expressed at relatively high levels in periportal hepatocytes whereas others are highly expressed in pericentral hepatocytes (Fig. S7A). Nevertheless, the differential expression across the liver lobule also decreased in general during the transition between the fed, fasted, and starvation states. Select canonical HIPPO/YAP target genes as well as *Yap1* are shown in Figure S7B. Similarly, 14 annotated Sonic hedgehog (SHH) signaling target genes are shown in the heatmap of Figure S7C, and selected marker genes, including *Shh*, are shown as bar graphs (Fig. S7D). As observed with both the WNT/β-catenin and HIPPO/YAP pathways, several mRNAs involved in SHH signaling display a loss of zonation during starvation (*Shh* and *Ptch1* but not *Sufu*). Consistent with these observed reductions in WNT/β-catenin target gene expression, Principal Component Analysis (PCA) showed that the overall changes in periportal gene expression are relatively small compared to the changes in pericentral gene expression (Fig. S8).

Moreover, as indicated with the arrow, as the hepatocytes transition from the fed to fasted and then starvation state, the pericentral hepatocytes modified their gene expression to resemble periportal hepatocytes. Taken together, these data are consistent with the COMPASS metabolic flux analysis (Fig. 2), suggesting that the increase in periportal-like hepatocyte gene expression in the pericentral hepatocytes during starvation is consistent with a general reduction in hepatic zonation as defined by mRNA expression.

### Reduced beta-catenin signaling redistributes expression of glutamine synthetase and glutaminase to increase pericentral hepatocytes glutamine-dependent gluconeogenesis

Several studies have reported that the *Gls2* gene (glutaminase) (downregulated by WNT) and *Glul* gene (glutamine synthetase) (upregulated by WNT) are canonical β-catenin liver target genes^50,51,42,52,53^, which is consistent with the observed increase in pericentral *Gls2* mRNA and decrease in pericentral *Glul* mRNA in the fasted/starvation states compared to the fed state (Fig. 7A). This was further confirmed by immunoblotting of glutaminase, where we saw an increase in the glutaminase protein in the isolated pericentral hepatocytes in the starved state (Fig. 7B). In contrast to glutaminase, glutamine synthetase (GS) immunofluorescence of fixed liver sections in the fed and starved (24h) state showed less glutamine synthetase protein expression in the starved state (Fig. 7C and D). The difference in intensity of GS expression could result from GS being restricted to only a few hepatocytes layers in the starved state.

**Fig. 7.**
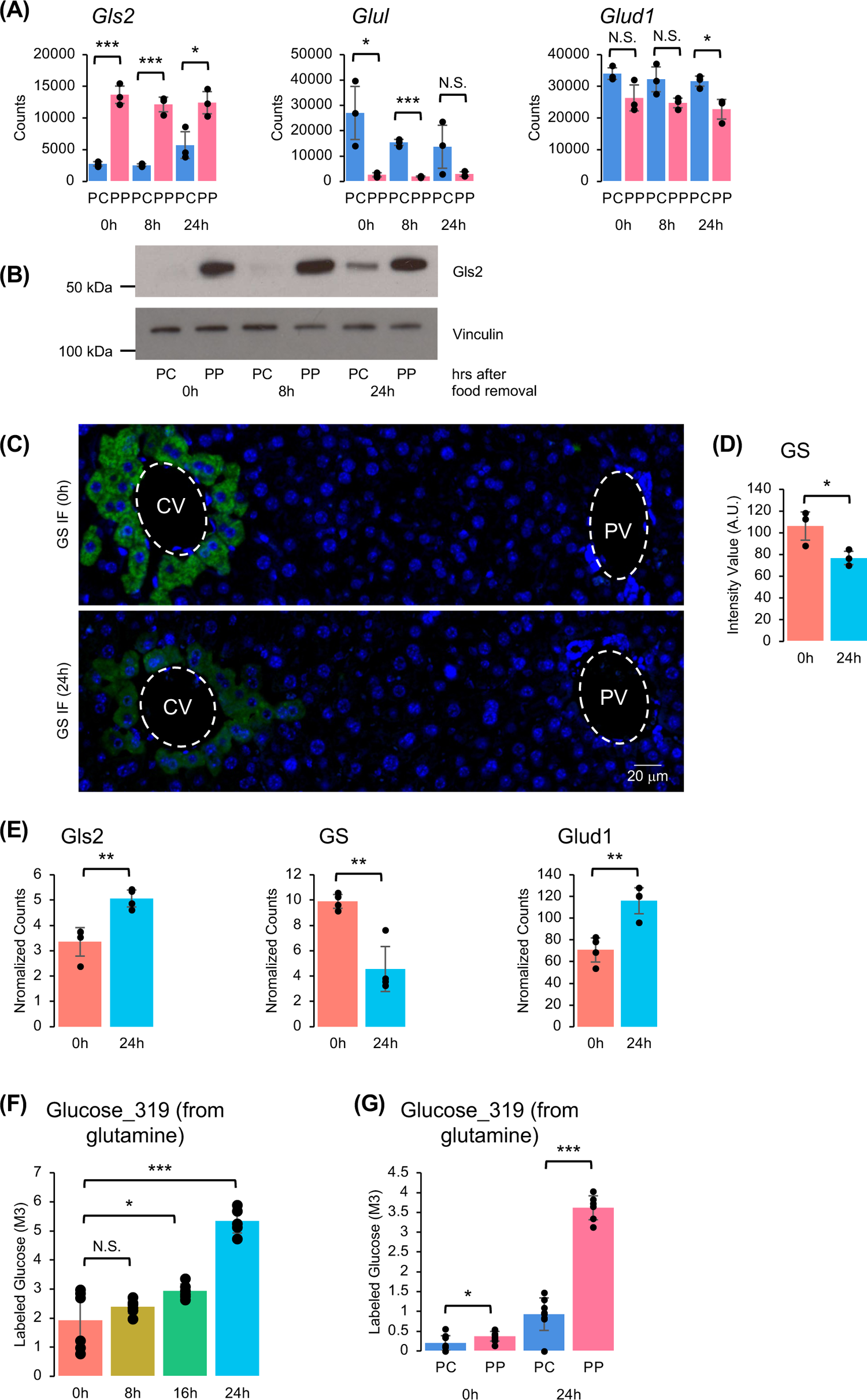
Fasting decreases Wnt signaling to increase glutamate synthesis. **(A)** RNA-seq data (counts) of *Gls2*, *Glul*, and *Glud1* from pericentral and periportal isolated hepatocytes. N=3 mice per time point/condition. **(B)** Representative immunoblotting of GLS2 and Vinculin from pericentral and periportal isolated hepatocytes. **(C)** Representative immunofluorescence images of GS from fed (0h) and fast24h (24h). **(D)** Quantification of GS immunofluorescence (Y-axis represents mean pixel intensity. Dots represent N=3 mice per time point.) **(E)** Proteomics (normalized counts) of GLS2, GS, and GLUD1 from total hepatocytes (N=1 mice per time point, dots represent technical replicates). **(F)** Glucose_319 measurement from [^13^C_5_] glutamine within each time point from total hepatocytes. X-axis represents time points; Y-axis represents labeled glucose (M3) (dots represent N=2 mice per time point/condition combined with technical replicates). ANOVA followed by Tukey’s multiple comparisons test was performed to compare 0h with different time points. **(G)** Glucose_319 measurement from [^13^C_5_] glutamine within each time point from pericentral and periportal isolated hepatocytes. X-axis represents time points; Y-axis represents labeled glucose (M3) (dots represent N=2-3 mice per time point/condition combined with technical replicates).

This observation is consistent with a recent report that hepatocyte glucagon receptor knockout mice (equivalent to the fed state) display an expansion of *Glul* expression from 1 to 2 hepatocyte layers around the central vein^52^. On the other hand, glutamate dehydrogenase (*Glud1*) mRNA does not display any significant zonation of expression and was unaffected by nutrient status (Fig. 7A). Proteomics analysis further confirmed the increase in glutaminase following 24h of food deprivation, and while no difference in *Glud1* mRNA was observed, proteomics identified that Glud1 protein nonetheless increased with fasting (Fig. 7E). Proteomics analysis showed a decrease in glutamine synthetase (GS) protein as well (Fig. 7E).

The reciprocal change in glutaminase and glutamine synthetase suggested that there would be a greater utilization of glutamine by the pericentral hepatocytes in starved state. To test this, we first examined the effect of fasting/starvation on glutamine-dependent gluconeogenesis using [^13^C_5_]-glutamine conversion to m3-glucose in total primary isolated hepatocytes (Fig. 7F). As observed for 2,3[^13^C_2_]-pyruvate, fasting for 8 and 16h resulted in only a marginal increase in glutamine incorporation into glucose.

However, at 24h of fasting there was a clear increase (∼2.5-fold) in glucose synthesis from glutamine. We then compared [^13^C_5_]-glutamine conversion to m3-glucose between pericentral and periportal hepatocytes. As expected, in the fully fed state the conversion of glutamine to glucose was relatively low with only a marginal difference between the pericentral and periportal hepatocytes (Fig. 7G). In contrast, glucose formation increased approximately 5-fold in the pericentral and approximately 10-fold in the periportal hepatocytes. The increase in pericentral glucose production from glutamine in the starved state is consistent with the zonation redistribution of glutamine synthetase and glutaminase.

## DISCUSSION

Previous studies in the 1980’s using isolated periportal and pericentral hepatocytes observed an approximate 2-fold difference in gluconeogenic activity between periportal and pericentral hepatocytes^19,20,21^. Similarly, other aspects of glucose/lipid metabolism were found to exhibit liver zonation differences, including glycolysis, de novo lipogenesis and fatty acid oxidation^15,16,17,22,23,24^. Although these were landmark studies, they were highly limited and did not address the temporal changes or molecular basis for these differences. As such, we established a well-defined protocol that synchronized the extent and time of feeding to delineate the transitions into the fasted and starvation states in mice.

Using single cell technologies (single cell RNA-seq, quantitative targeted single cell RNA-seq and single-molecule fluorescence in situ hybridization), our data clearly demonstrates that lipogenic gene expression is relatively equally distributed between periportal and pericentral hepatocytes and is quickly suppressed when food intake ceases. In contrast, gluconeogenic gene expression upon the induction of fasting is primarily localized to the periportal hepatocytes but as the time of food deprivation increases into the starvation state, gluconeogenic gene expression expands across the liver lobule from the periportal to the pericentral hepatocytes. As expected, since *Pck1* protein (PEPCK) has a half-life of six hours^54,55^, the zonation changes in *Pck1* mRNA paralleled the changes in PEPCK protein. However, the induction of gluconeogenic activity was substantially slower than either the changes in *Pck1* mRNA or PEPCK protein, clearly showing that changes in *Pck1* mRNA cannot be used as surrogate to make any conclusions about gluconeogenic activity. More importantly, the hepatocytes across the lobule show a high degree of plasticity in their ability to express gluconeogenic genes and gluconeogenic activity that is directly dependent upon the specific time of nutrient deprivation. This underscores the critical importance of using well-defined feeding and fasting times, as the expression and function of gluconeogenesis is highly dependent upon the specific conditions employed, and as such, may account for many of the divergent findings in studies trying to define the basis of hepatic insulin resistance and glucose production.

There are many mechanisms that could account for the metabolic plasticity across the liver lobule including changes in the oxygen gradient, nutrient flow, glucagon/insulin gradients, intercellular signals (ie: gap junctions) and neural signaling. Our RNA-seq data demonstrated a progressive reduction in the canonical WNT signaling gradient as seen through the reduced expression of β-catenin target genes as the mice transition into the fasted and then starvation state, in parallel to the increased expression of *Pck1* and *G6pc* mRNA in the pericentral hepatocytes. Moreover, the pericentral hepatocytes temporally acquire a more periportal hepatocyte phenotype, at least at the level of gene expression. Sarkar et al have reported that intermittent fasting interacts with WNT signaling, which is consistent with our findings^56^. Previous studies have demonstrated that the WNT signaling gradient is established at the time of weaning and is a necessary regulator of hepatocyte lobule zonation^57,58, 59^. Thus, we speculated that high β-catenin activity in the pericentral hepatocytes might suppress gluconeogenic gene expression and this suppression is released during fasting as β-catenin signaling declines. In this regard, β-catenin was reported to function as a co-activator of FoxO1 to enhance *Pck1* gene expression and gluconeogenesis^60^. Similarly, Liu et al reported that β-catenin can transactivate *Foxo1* in the fasted state to enhance *Pck1* gene expression by reducing β-catenin interaction with the canonical WNT signaling to TCF^61^. Since these previous studies examined total hepatocytes, the most likely explanation for these differences is the dependence of β-catenin /FoxO1 transcriptional activation in pericentral versus periportal hepatocytes.

In any case, although our study does not address the mechanism by which β-catenin co-transcriptional activation of TCF gene targets is reduced while FoxO1 targets are increased, it does demonstrate the functional consequence, at least one of which is to increase pericentral PEPCK and glutaminase while decreasing glutamine synthetase protein levels. The net effect would be to enhance the conversion of glutamine to glutamate and subsequently increase the formation of oxalacetate in the mitochondria pericentral hepatocytes. In turn, oxalacetate exit from the mitochondria by the glutamate-aspartate translocase and the cytoplasmic oxalacetate can then serve as substrate for PEPCK to further drive gluconeogenesis^62,63^. This mechanism accounts for, at least in part, the basis for increased pericentral gluconeogenesis that occurs in the starved state. Further studies will be needed to determine whether the reduction in pericentral hepatocyte β-catenin signaling in the starved state results from a direct effect on β-catenin itself or upstream signaling events such as WNT2b, WNT9 and/or Rspo1 reported to be necessary activators of pericentral β-catenin co-transcriptional activation^45,46,64^.

## METHODS

### Animals

All studies followed the guidelines that were approved and in agreement with the Albert Einstein College of Medicine Institutional Animal Care and Use Committee.

All experiments were performed on 10- to 14-week-old C57BL/6J WT male mice from The Jackson Laboratory (#000664). Mice had free access to low fat (4.5%) laboratory mouse chow (LabDiet #5053) and water unless otherwise noted. All mice were housed in groups and under a facility equipped with a 12h light/dark cycle. As previously described^26^, prior to fasting and feeding experiments, mice were trained for 3 days by removing food at 5:00 and feeding at 9:00 the next day (16h fasting overnight and 8h feeding daytime). On the day of experiment, mice were provided food with drinking water that contained 20% of sucrose (w/v) (sucrose water was prepared and provided to mice on the day of experiment and not during the fasting/feeding training period). The fed (0h) mice were sacrificed at 13:00 equal to 4h of feeding (0h). For fasted mice, the food and sucrose water were taken out at 13:00, then sacrificed at 17:00 for fast 4h (4h), 21:00 for fast 8h (8h), 5:00 the next day for fast16h (16h), 13:00 the next day for fast 24h (24h), and 19:00 the next day for fast 30h (30h).

### Hepatocyte isolation

Liver perfusion was done under anesthesia, and the procedure combined multiple protocols previously described by literature^37,65^. Briefly, for total hepatocyte isolation, a catheter (TERUMO, #SR-OX2419CA) was placed in the portal vein (PV) and the lower inferior vena cava (IVC) was cut open for drainage of fluid. For pericentral/periportal hepatocyte isolation, one catheter was placed in the PV while the other catheter was placed in the IVC. The catheter was then connected to a perfusion line (PV for total and periportal, IVC for pericentral), and the liver was perfused with pre-warmed 1xPBS. While the liver was perfused with 1xPBS, an artery clip was placed on the upper IVC. The perfusion line was then turned off and switched to pre-warmed EBS buffer containing liberase (Sigma-Aldrich, #5401127001). For pericentral/periportal hepatocyte isolation, 833 μL of 5-10mM digitonin (Sigma-Aldrich, #D141-500MG) in a 1 mL syringe was injected from the unconnected catheter (IVC for periportal, PV for pericentral), then the unconnected catheter was removed from the vein and the vein was cut. The perfusion was restarted immediately until 15 mL of EBS buffer containing liberase was consumed. The liver was cut from the mouse and placed on a pre-chilled 10cm dish containing 20 mL of DMEM (Corning, #90-113-PB). Dissociated liver cells were transferred to a 50 mL conical tube with a 100 μm cell strainer placed over the top (Fisher Scientific, #22363549). Liver cells were centrifuged at 40 x *g* for 5 min at 4 °C. The pellet (enriched with hepatocytes) was resuspended in 10 mL DMEM. To discard dead hepatocytes, 10 ml of Percoll solution (9 ml Percoll (Sigma-Aldrich, #P4937-500ML) mixed with 1 ml 10× PBS (Invitrogen, #AM9625)) was added to the cells and centrifuged at 80 x *g* for 10 min at 4 °C. The supernatant containing dead cells was aspirated and the pellet was resuspended in 10 ml DMEM, then centrifuged at 40 x *g* for 5 min at 4 °C. The supernatant was aspirated, and the pellet resuspend in the preferred amount of DMEM for quantification and assessment. For stable isotope experiments, DMEM was replaced with KH Buffer.

### Single-cell RNA sequencing (scRNA-seq)

#### Sample preparation

Isolated total hepatocytes were placed into the 10x Genomics Chromium Controller. The cDNA library from each time point (0h, 4h, 8h, 16h, 24h, and 30h, N=1 each) was prepared independently with Chromium Next GEM Single Cell 3’ GEM Kit v3 according to the company’s manual. The sequencing was done with sequencing by synthesis (Illumina).

#### Data processing (bioinformatics)

The raw FASTQ files are mapped to single cells and gene expression levels were reconstructed using Cell Ranger (v6.0.1, 10X Genomics) with a default parameter. Seurat (v4.0.5) standard workflow as well as Loupe Browser (v6.4.1) were used for data analysis and clustering. Zonation markers (such as *Cyp2e1*, *Gulo*, *Cyp2f2*, *Hsd17b13*) that were previously reported^27,66,28^ were used to identify pericentral and periportal hepatocytes. Because standard scRNA-seq is not quantitative for low-abundance transcripts^30^ and the expression levels of zonation markers change upon fasting/feeding, rather than using thresholds for clustering, polygonal selection with Loupe Browser was performed.

#### COMPASS cellular metabolic state analysis

Pericentral and periportal clusters from 0h, 8h, and 24h were compared using the COMPASS method established by Wagner et al and Cohen’s D analysis was performed based on their method^29^. The Compass python implementation (https://github.com/YosefLab/Compass) was used.

#### Targeted gene expression analysis (10x Genomics)

The cDNA library from standard scRNA-seq went through the targeted gene expression protocol (Target Hybridization Kit, PN-1000248) and NovaSeq sequencing. Raw FASTQ files were mapped to single cells and 61 gene expression levels were reconstructed using Cell Ranger (v6.0.1, 10X Genomics) in Targeted Gene Expression mode. To search for patterns in the time profiles of the regulated gene expression, we applied fuzzy c-means (FCM) clustering which was extensively used to elucidate signaling dynamics from phosphoproteome data and applied this method to the targeted scRNA-seq dataset^31^. We iteratively explored combinations of cluster sizes and parameter ***m*** and found optimal partitioning with c = 8 and m = 1.

### Single-molecule Fluorescence *in situ* Hybridization (smFISH)

#### Sample preparation

Livers collected at specific time points were rinsed with 1xPBS, immediately fixed in cold 4% paraformaldehyde (Electron Microscopy Sciences, #15710-S) in 1xPBS (v/v) for 3h at 4 °C with gentle rolling, then fixed overnight in cold paraformaldehyde + 30% sucrose (w/v) at 4 °C with gentle rolling. The liver was taken out of paraformaldehyde the next day, rinsed with 1xPBS, then embedded in pre-chilled OCT (SAKURA, #4583) in a cryomold (SAKURA, #4557), and put in cold 2-methylbutane in dry-ice. Tissue blocks were kept at −80 °C until sectioning. Sections were 8-10 μm thick and were sectioned in a −20 °C/-20 °C cryostat.

#### smFISH and imaging

Frozen liver sections were fixed with 4% (v/v) paraformaldehyde in 1xPBS for 15 min followed by a brief wash with 1xPBS. Sections were then incubated in cold 70% ethanol for at least 2h before hybridization. After treating with proteinase K (Ambion, #AM2546) in 2x SSC buffer (Invitrogen, #15557-044) for 15 min at 50 °C, the sections were placed in pre-hybridization buffer (20% formamide (Fisher Scientific, #AC205821000) (v/v) in 2x SSC buffer) for 1h, followed by overnight incubation at 37 °C in a humified chamber with 50-100 nM Stellaris RNA FISH probes (LGC Biosearch Technologies) labeled with Quasar 670 or Quasar 570 fluorescent dye and complementary to the coding sequence of each gene (*Pck1*, *Fasn*, *Glul*), in hybridization buffer (10% dextrane sulfate, 20% formamide, 1 mg/ml E.coli tRNA, 2mM VRC, 0.2 mg/ml BSA in 2x SSC) as previously described^67^. The following day, sections were washed in 20% formamide in 2x SSC/RNase-free water twice for 30 min at 37 °C. After washing, DNA was counterstained with DAPI and mounted with Prolong Diamond antifade on the slides (Invitrogen, #P36962). All images were taken with an upright, wide-field Olympus BX-63 Microscope equipped with a Super Apochromatic objective lens (60×/1.35 N.A., Pixel size (x,y) = 107.5 nm, z-step = 300nm). MetaMorph software (Molecular Devices) was used for controlling microscope automation and image acquisition. Pericentral hepatocytes were identified based on Glul FISH, while periportal hepatocytes were identified based on the morphology of the bile duct (DAPI). Liver lobule images were taken by taking sequential images and stitching them on FIJI (v2.14.0). For transcription site analysis, images were analyzed using FISH-quant v2 (FQ ImJoy plugin: v0.0.16, FQ interface: v0.0.11, big-fish: v0.6.0)^32^. Non-hepatocyte nuclei were excluded during the quantification process. smFISH probe sequences were based on previous studies by Halpern et al and are available in table 2^35,66^.

**Table 2.**
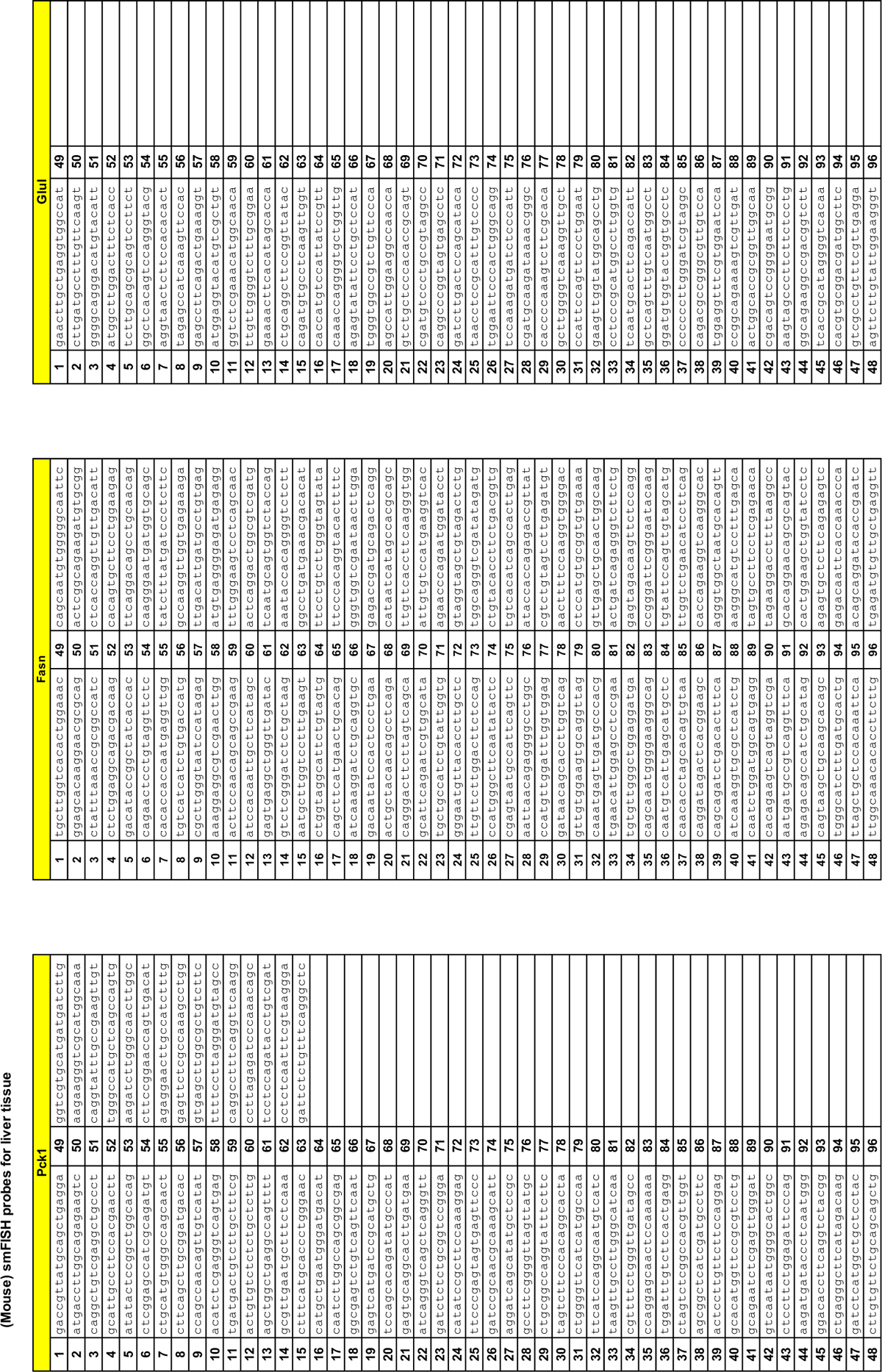
smFISH probe sequences.

### RNA isolation and quantitative PCR (RT-qPCR)

Total RNA isolation from isolated hepatocytes and mouse liver was done with RNeasy mini kit (Qiagen, #74106) combined with TRIzol (Invitrogen, #15596026) according to the manufacture’s protocol. cDNA strand synthesis and RT-qPCR were done with SuperScript IV VILO cDNA synthesis kit (Invitrogen, #11756050) and PowerUp SYBR Green Master Mix (Applied Biosystems, #A25778). RT-qPCR reactions, measurements and quantification were performed on QuantStudio3/QuantStudio6 (Applied Biosystems) according to the manufacture’s protocol. Samples were normalized to *Ppib* gene to determine relative mRNA levels. Primer sequences used in all experiments are listed in table 3^26^.

**Table 3.**
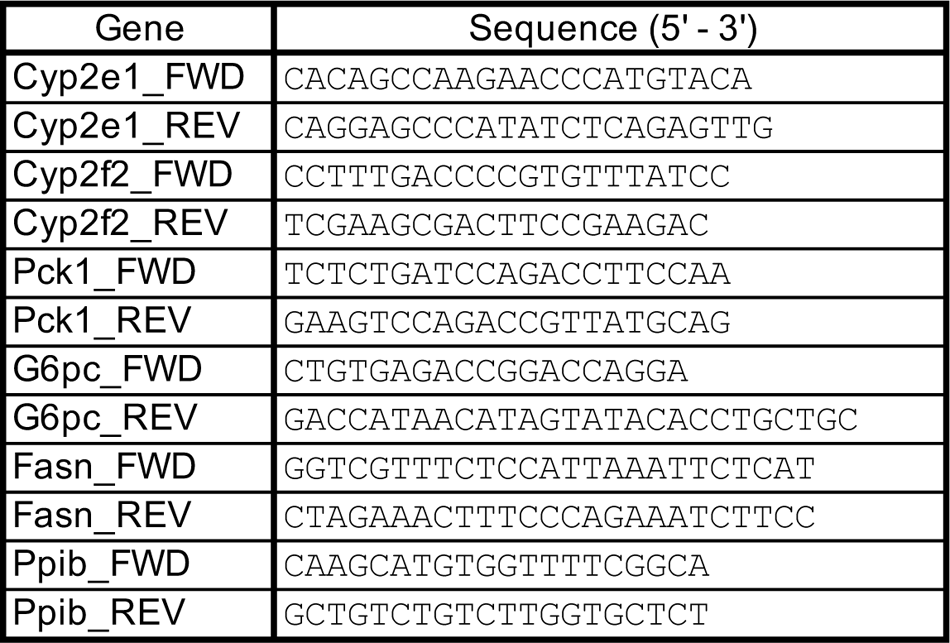
RT-qPCR primer sequences.

### Bulk RNA sequencing

After assessing pericentral/periportal isolated total RNA samples with NanoDrop (ThermoFisher Scientific), 2 μg of the RNA were sent to Azenta Life Sciences for genome-wide mRNA sequencing using NEBNext Ultra RNA Library Prep Kit (New England Biolabs, Ipswich, MA) for NovaSeq (Illumina) sequencing by synthesis. All downstream analysis was performed using R (v4.1.3). Transcript-level raw counts from Kallisto (v0.46.0) were imported into R using the Bioconductor package tximport (v1.20.0), and the expression levels of each gene were estimated. Differential expression analysis between the two groups was performed using the DESeq2 (v1.32.0), which generated log2(Lfold change) and adjusted *P*-value.

### Immunoblotting

Commercially available primary antibodies were purchased for the detection of PEPCK (Abcam, #ab70358), Cyp2e1 (Invitrogen, #PA5-52652), GS (Invitrogen, #MA5-27749), E-cadherin (Cell Signaling Technology, #3195), GLS2 (Abcam, #ab113509), and Vinculin (Santa Cruz, # sc-73614). Isolated pericentral/periportal hepatocytes were rocked for 10 min at 4 °C with RIPA lysis buffer (ThermoFisher Scientific, #89900) with protease and phosphatase inhibitors (Sigma-Aldrich, #S8830-20TAB, #P5726). After centrifugation, the total protein in the supernatant was quantified by BCA assay (ThermoFisher Scientific, #23225), and the samples were normalized by total protein content. The samples were then resuspended in Laemmli buffer (BIO-RAD, #1610747) and heated at 100 °C for 5 min. Samples were separated by SDS polyacrylamide gel electrophoresis (BIO-RAD, #4561094) and electrophoretically transferred to polyvinylidene difluoride (PVDF) membranes. The membrane was blocked with either 1% BSA (w/v) (Sigma-Aldrich, #A7906-50G) or 5% Milk (w/v) (Santa Cruz, #sc-2324), then incubated with the primary antibody in primary antibody solution. The membrane was then washed in TBS with Tween20 (TBST) and incubated with horseradish peroxidase-conjugated secondary antibody in 5% milk. The membrane was washed with TBST and visualized either by enhanced chemiluminescence (ECL) (Cytiva, #RPN2232). ImageJ was used for quantification.

### Stable Isotope flux analysis

After the final wash with KH Buffer, 100,000 cells of the isolated total, pericentral or periportal hepatocytes were incubated with 1mM 2,3 [ ^13^C_2_] sodium pyruvate (or [^13^C_3_] sodium pyruvate or unlabeled sodium pyruvate), 1mM sodium lactate, 2mM ^13^C_5_ L-glutamine (or unlabeled L-glutamine), 1mM BSA-conjugated Palmitic acid in KH Buffer at 37 °C for 1h in a shaking water bath. After incubation, the cells were centrifuged at 100 x *g* for 5 min at 4 °C. The media was collected to a screw vial and snap frozen in liquid nitrogen. Following thawing on ice, 200 μl of the media was added to 800 μl of methanol. After a brief vortex, the media was centrifuged at 13,100 x *g* for 10 min at 4 °C. The supernatant was transferred to a glass vial and dried under gentle nitrogen flow. The samples were then subjected to a 2-step derivatization with 50 µl of methoxyamine hydrochloride (MOA, 15 mg/mL in pyridine) at 30 °C for 90 minutes, and then incubated with 50 µl of N,O-Bis(trimethylsilyl)trifluoroacetamide (BSTFA, containing 1% TMCS) at 70 °C for 60 min. The samples were analyzed by gas chromatography mass spectrometry (GC (8890)-MS (5977B), Agilent). Citrate_465 was used to calculate PC/PDH ratio. The data was first processed with MassHunter software (Agilent). Isotope enrichment calculation (background correction) was performed as described by Jennings and Matthews^68^.

### Immunofluorescence

Livers from fed and fasted mice were fixed as described (see smFISH methods section). After fixation, samples underwent ethanol dehydration, xylene clearing, paraffin embedding, and sectioning at 5-micron thickness. FFPE sections were baked in a 65°C oven for 2h and then transferred to a Leica Bond RX automated stainer for sequential multifluorescent staining. Briefly, each slide was baked for an additional 30 min, dewaxed with Bond Dewax solution (Leica Biosystems, #AR9222) and subjected to epitope retrieval with Bond ER solution 2 (Leica Biosystems, #AR9640). Antibody labeling was then accomplished by 4 sequential rounds of primary antibody incubation, secondary antibody incubation, fluorescent tyramide incubation and antibody stripping. Mouse IgG anti-glutamine synthetase (Invitrogen, #MA5-27749) was diluted in Bond Wash (Leica Biosystems, #AR9590) containing 1% BSA at 2.5 µg/ml. Opal anti-mouse and rabbit HRP secondary antibody (Akoya Biosciences, #ARH3001KT) was used without dilution. Fluorescent tyramide CF488A (Biotium, #92171) dissolved in DMSO was diluted in 1x Plus Automation Amplification Diluent (Akoya Biosciences, #FP1609) at 6.7 µM. Hoechst 33342 (Cell Signaling, #4082) was diluted to 2.5 µM in Bond Wash. Slides were scanned with a PhenoImager HT operating with Vectra Polaris software version 1.0.13 (Akoya Biosciences) using the multispectral whole slide scan function and a 20x objective.

Images were viewed using QuPath (v0.4.3^69^). GS Quantification was performed on 6 whole-slide scans in QuPath (3 fed, 3 fasted) using the “Pixel classification” tool (Gaussian prefilter, Smoothing sigma = 0, Threshold = 50) and the “Add intensity features” tool (Resolution = 0.5 µM).

### Proteomics

Hepatocytes were isolated from fed and fasted mice as described above and processed in isolation buffers (PBS, EBS, and KH buffers) containing 5 mM glucose. Hepatocytes were aliquoted at 1 million cells into four tubes, washed with 150 mM ammonium acetate in water, and snap frozen in liquid nitrogen. Cell pellets were homogenized with a probe sonicator in buffer containing 5% SDS, 5 mM DTT and 50 mM ammonium bicarbonate (pH = 8) and incubated for 1h at room temperature.

Samples were then alkylated with 20 mM iodoacetamide in the dark for 30 min. Afterward, phosphoric acid was added to a final concentration of 1.2%. Samples were diluted in six volumes of binding buffer (90% methanol and 10 mM ammonium bicarbonate, pH 8.0). After gentle mixing, the protein solution was loaded into an S-trap filter (Protifi) and spun at 500 x *g* for 30 sec. The sample was washed twice with binding buffer. Finally, 1 µg of sequencing grade trypsin (Promega), diluted in 50 mM ammonium bicarbonate, was added into the S-trap filter and samples were digested at 37°C for 18 h. Peptides were eluted in three steps: (i) 40 µl of 50 mM ammonium bicarbonate, (ii) 40 µl of 0.1% TFA and (iii) 40 µl of 60% acetonitrile and 0.1% TFA. The peptide solution was pooled, spun at 1,000 g for 30 sec and dried in a vacuum centrifuge. Sample desalting and LC-MS/MS acquisition and analysis was performed as described elsewhere^70,71^.

### Glycogen quantification

Glycogen levels in total hepatocytes were measured by using the glycogen assay kit (Sigma-Aldrich, #MAK016-1KT) according to the manufacture’s protocol.

### Quantification and statistical analysis

Excel (v16.83) and Prism (v10.2) GraphPad Software were used for data processing, analysis, and graph productions in the experiments. All data are presented as the mean +/- standard deviation unless otherwise noted. Student’s *t*-test with unpaired two-tailed *p* values of \**p*<0.05, \*\**p*<0.01, and \*\*\**p*<0.001 was considered statistically significant when comparing two groups. (ns=not statistically significant).

## DATA AVAILABILITY

Raw fastq files were deposited to the National Center for Biotechnology Information Gene Expression Omnibus database under accession numbers: GSE263415, GSE263418, and GSE263419. Raw proteomics files will be available before publication.

## CODE AVAILABILITY

This study was conducted using only publicly available software; no custom code was used.

## ACKNOWLEDGEMENTS

This study was supported by grants (DK110063, DK020541, P30CA013330, S1OD030286) from the National Institutes of Health. We thank Shiori Okada for data analysis, Dr. Lawrence Leung for immunofluorescence, and Dr. David Reynolds for the scRNA-seq processing.

---

**Fig. S1.**
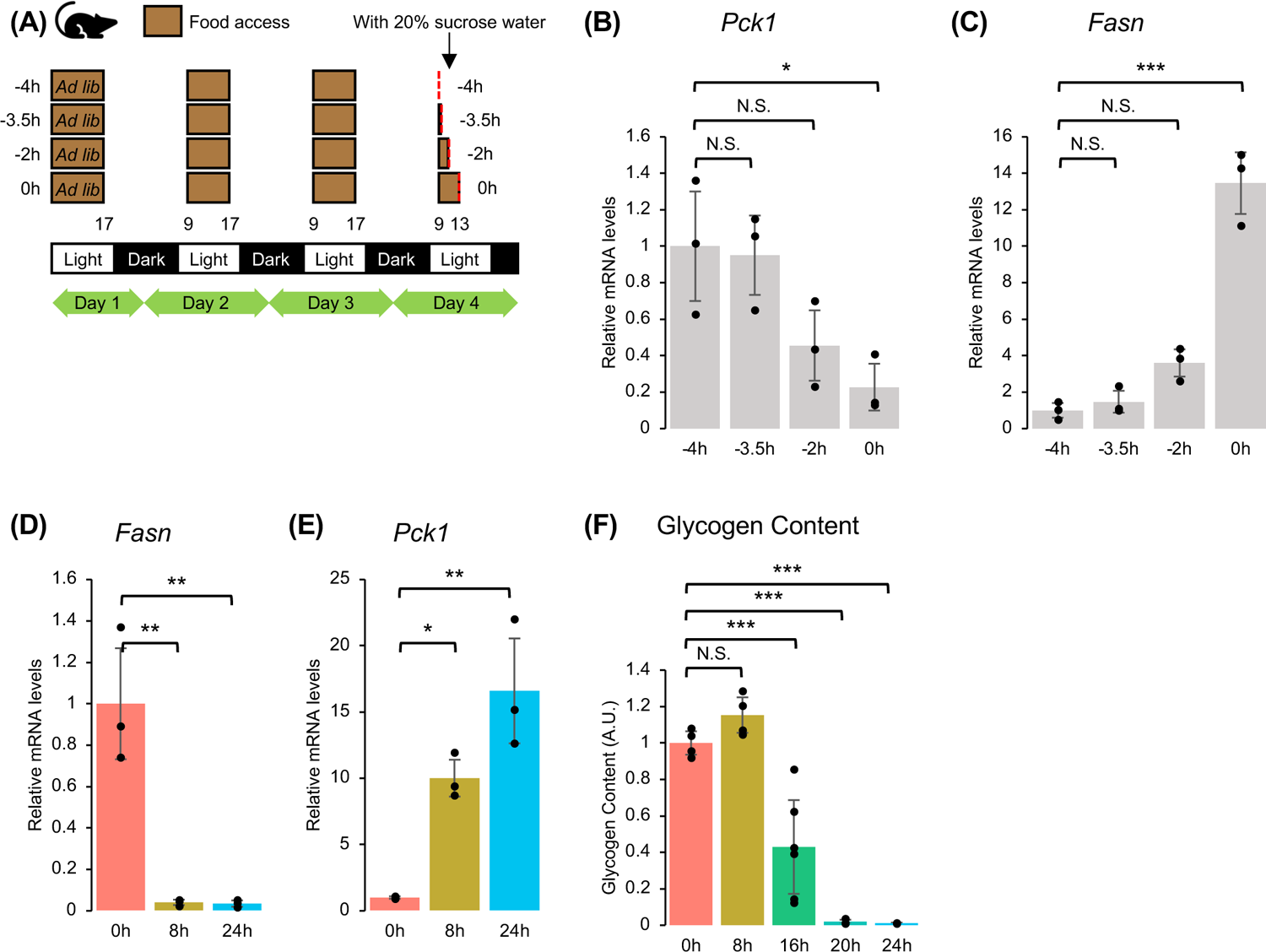
**(A)** Scheme of the feeding time course to determine the fully fed state. **(B, C)** RT-qPCR (relative mRNA levels) of *Pck1* **(B)** and *Fasn* **(C)** from total hepatocytes in the feeding time course. N=3 mice per time point/condition. ANOVA followed by Tukey’s multiple comparisons test was performed to compare −4h with different time points. **(D, E)** RT-qPCR (relative mRNA levels) of lipogenic (*Fasn*) **(D)** and gluconeogenic (*Pck1*) **(E)** genes from total hepatocytes in the fed (0h), fasted (8h) and starvation (24h) state. N=3 mice per time point/condition. ANOVA followed by Tukey’s multiple comparisons test was performed to compare 0h with different time points. **(F)** Glycogen content (A.U.) from total hepatocytes in each time point. Dots represent N=2-3 mice per time point/condition combined with technical replicates. ANOVA followed by Tukey’s multiple comparisons test was performed to compare 0h with different time points.

**Fig. S2.**
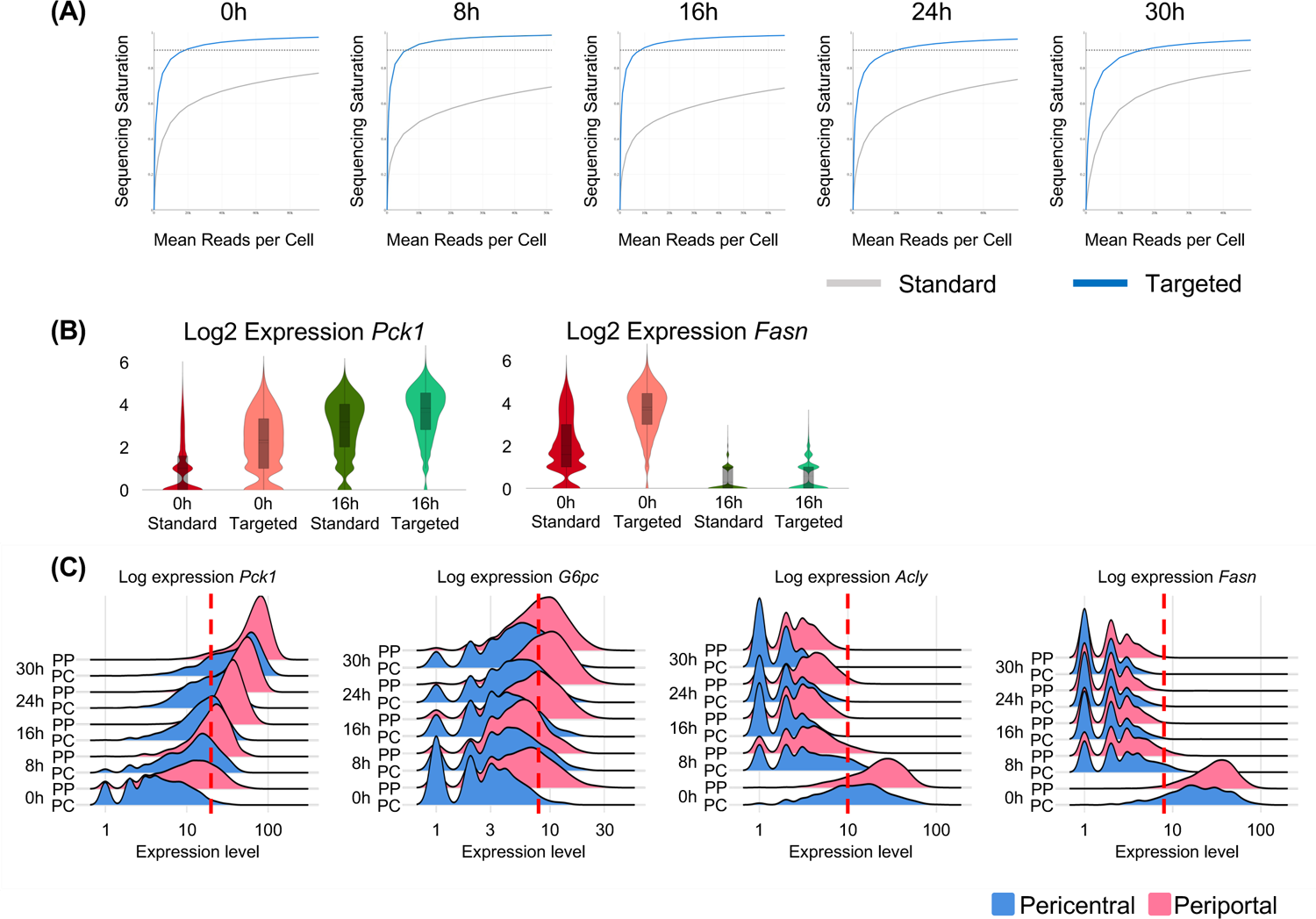
**(A)** Sequencing saturation plot of each library. X-axis represents mean reads per cell; Y-axis represents percentage of transcripts sequenced; dotted line is the approximate saturation point. **(B)** Violin plots of Log2 expression *Pck1* and *Fasn* comparing standard and targeted scRNA-seq in 0h and 16h. **(C)** Ridge plot analysis of gluconeogenic (*Pck1*, *G6pc*) and lipogenic (*Acly*, *Fasn*) genes from targeted scRNA-seq. X-axis represents log2 expression; Y-axis represents each time point/condition, red dotted line shows the threshold for cells positive of each gene.

**Fig. S3.**
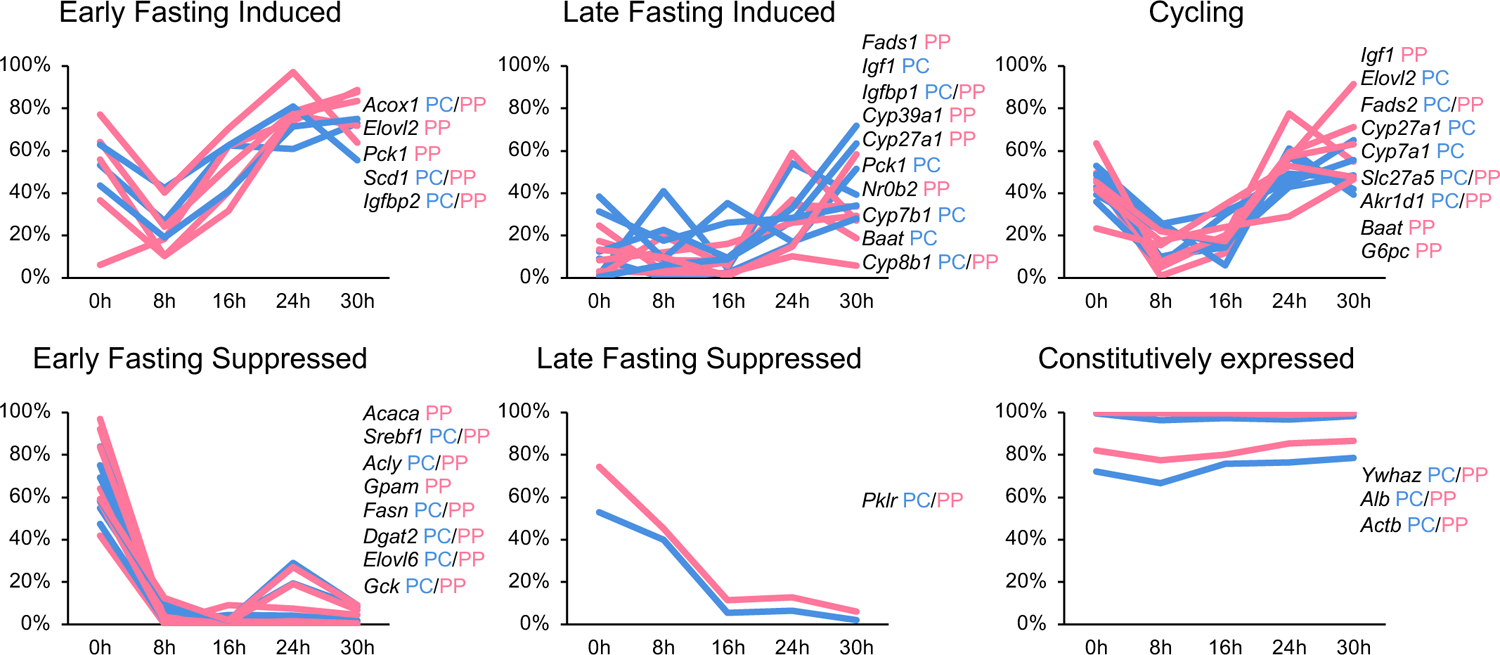
MFuzz^31^ analysis of the targeted scRNA-seq to cluster similar expression profiles. Once percentage of positive cells were defined based on ridge plot analysis (Fig. S2), each gene expressed in pericentral and periportal hepatocytes were assigned to 8 clusters; early fasting induced, late fasting induced, cycling, early fasting suppressed, late fasting suppressed, constitutively expressed, low expressed (not shown), and other (not shown). X-axis represents time points; Y-axis represents percentage of positive cells.

**Fig. S4.**
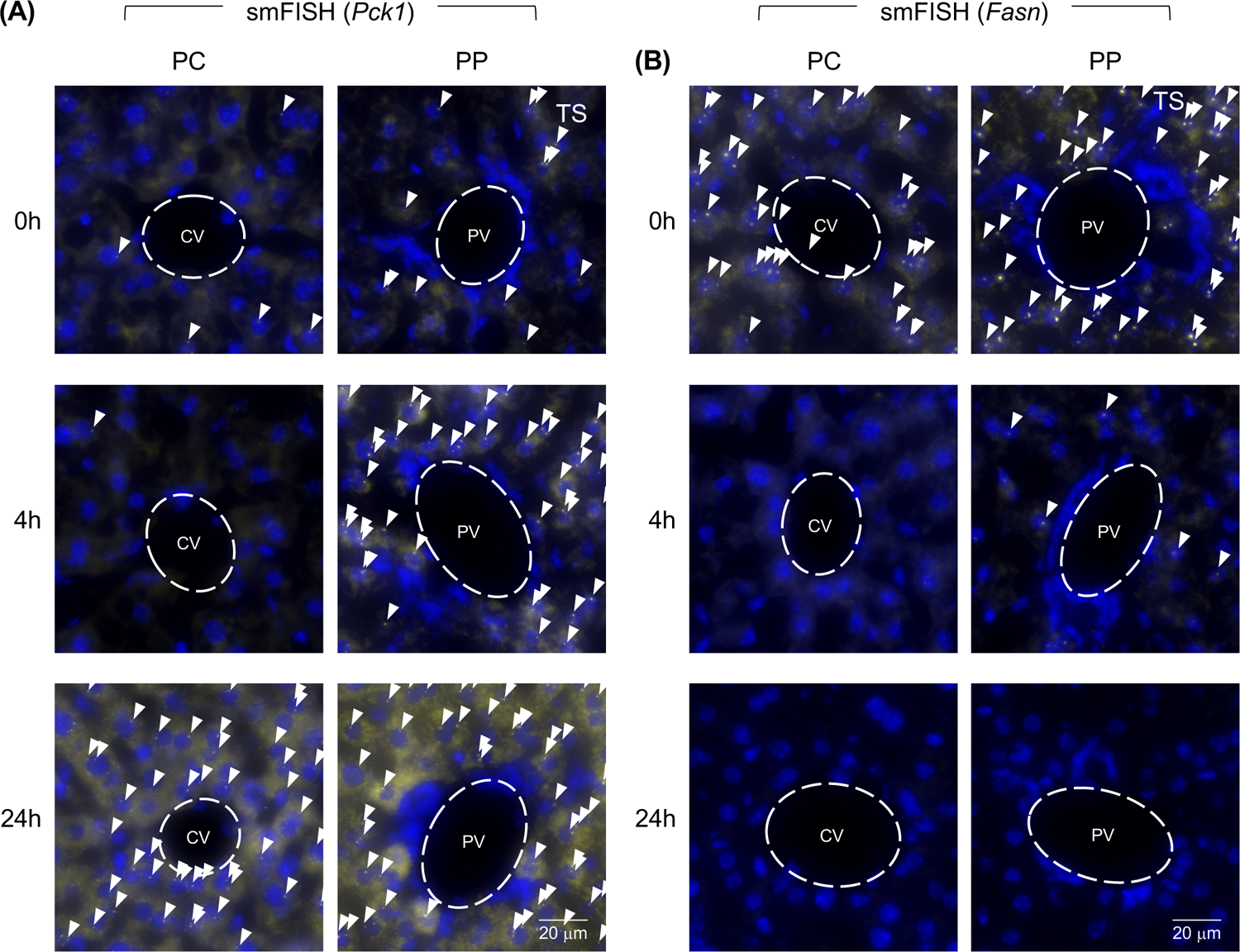
**(A, B)** Representative smFISH images of pericentral (PC) and periportal (PP) areas selected for quantification of TS (*Pck1* **(A)**, and *Fasn* **(B)**) from 0h, 4h, and 24h. TS are highlighted with arrows.

**Fig. S5.**
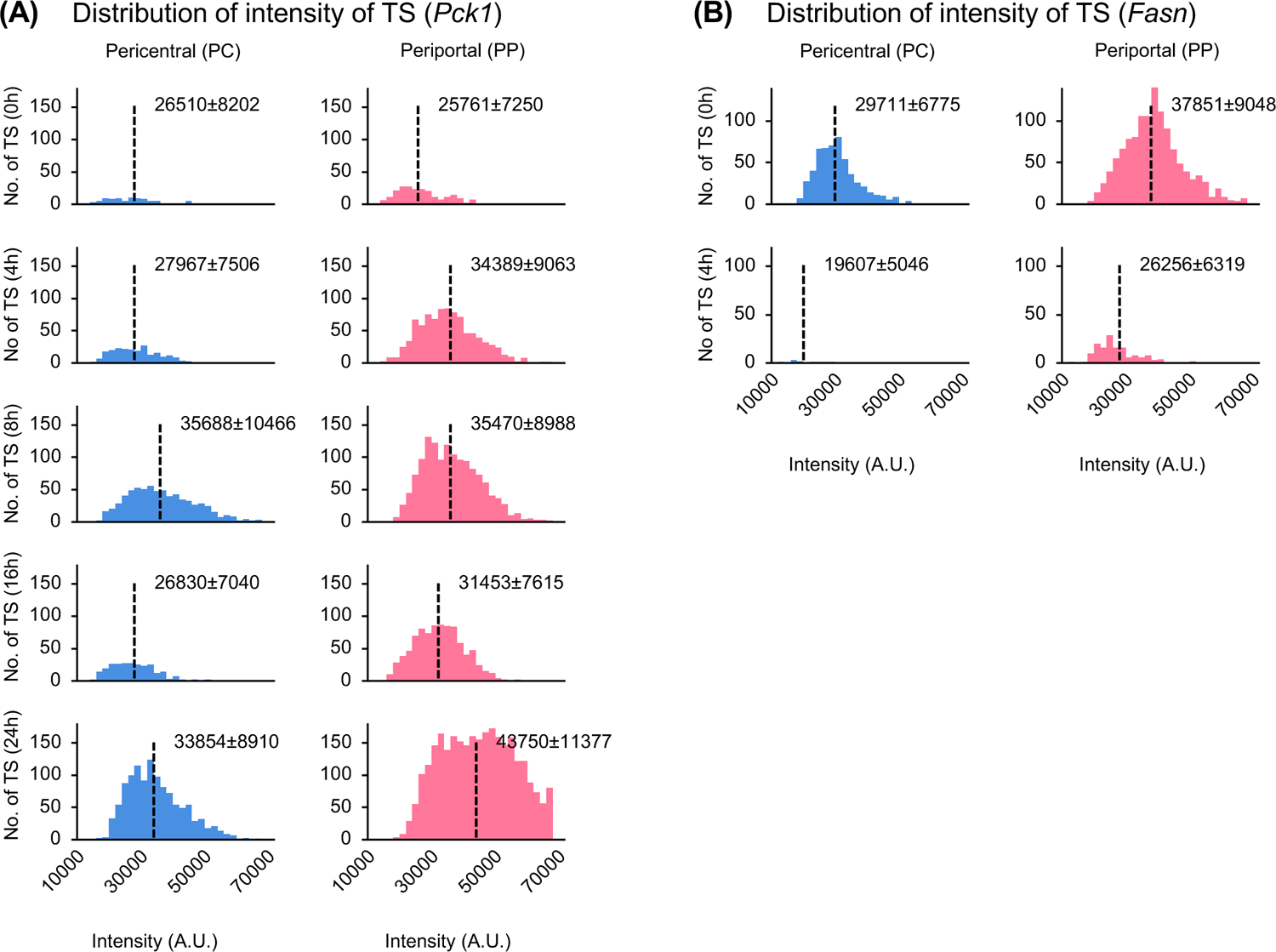
**(A, B)** Histogram based on the intensity of each TS (*Pck1* **(A)**, and *Fasn* **(B)**). X-axis represents the intensity of TS (A.U.); Y-axis represents the number of TS within each intensity. Dotted line shows the mean value of the intensity of TS with SD.

**Fig. S6.**
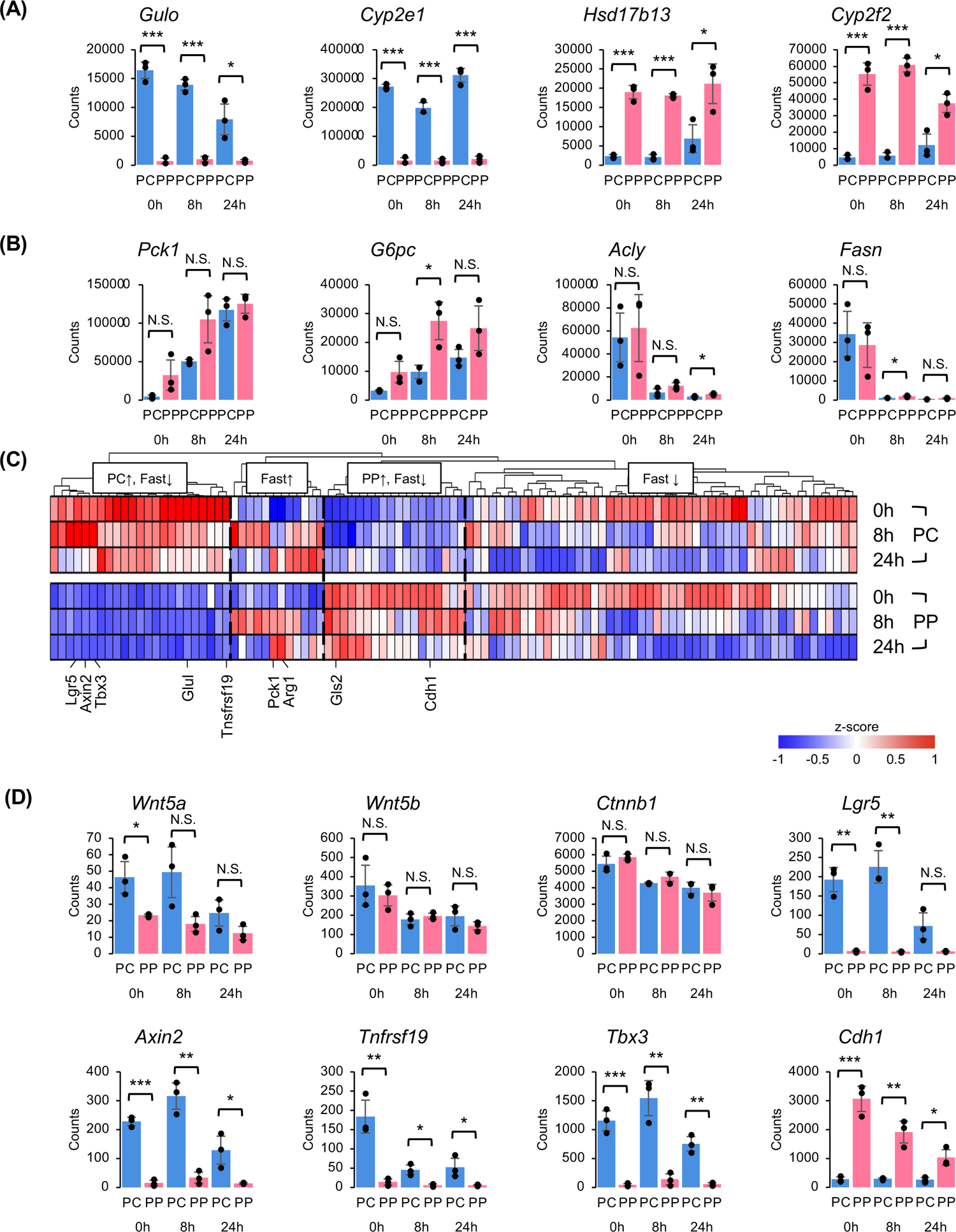
**(A, B)** RNA-seq data (counts) of pericentral zonation markers (*Gulo* and *Cyp2e1*) and periportal zonation markers (*Hsd17b13* and *Cyp2f2*) **(A)**, gluconeogenic genes (*Pck1*, *G6pc*) and lipogenic genes (*Acly*, *Fasn*) **(B)** from pericentral and periportal isolated hepatocytes. N=3 mice per time point/condition. **(C)** Heatmap of WNT target genes in RNA-seq from pericentral and periportal isolated hepatocytes. Average expression of N=3 mice from each time point/condition is shown. Coloring based on z-score. 4 main clusters show PC upregulated/fast reduced, fast induced, PP upregulated fast reduced, fast reduced **(D)** RNA-seq data (counts) of WNT signaling related genes from pericentral and periportal isolated hepatocytes. N=3 mice per time point/condition.

**Fig. S7.**
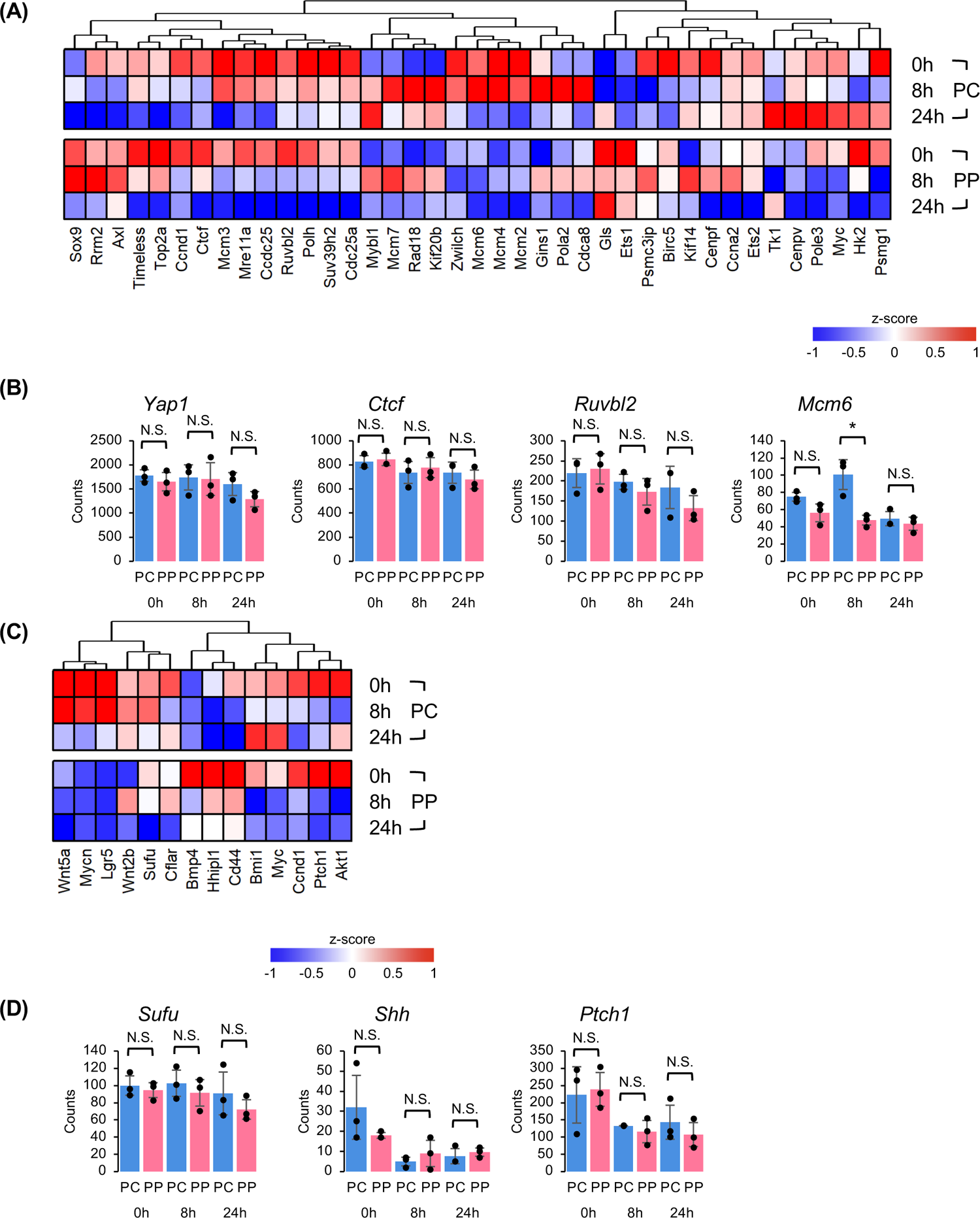
**(A)** Heatmap of Yap target genes in RNA-seq from pericentral and periportal isolated hepatocytes. Average expression of N=3 mice from each time point/condition is shown. Coloring based on z-score. **(B)** RNA-seq data (counts) of Yap related genes from pericentral and periportal isolated hepatocytes. N=3 mice per time point/condition. **(C)** Heatmap of Shh target genes in RNA-seq from pericentral and periportal isolated hepatocytes. Average expression of N=3 mice from each time point/condition is shown. Coloring based on z-score. **(D)** RNA-seq data (counts) of Shh related genes from pericentral and periportal isolated hepatocytes. N=3 mice per time point/condition.

**Fig. S8.**
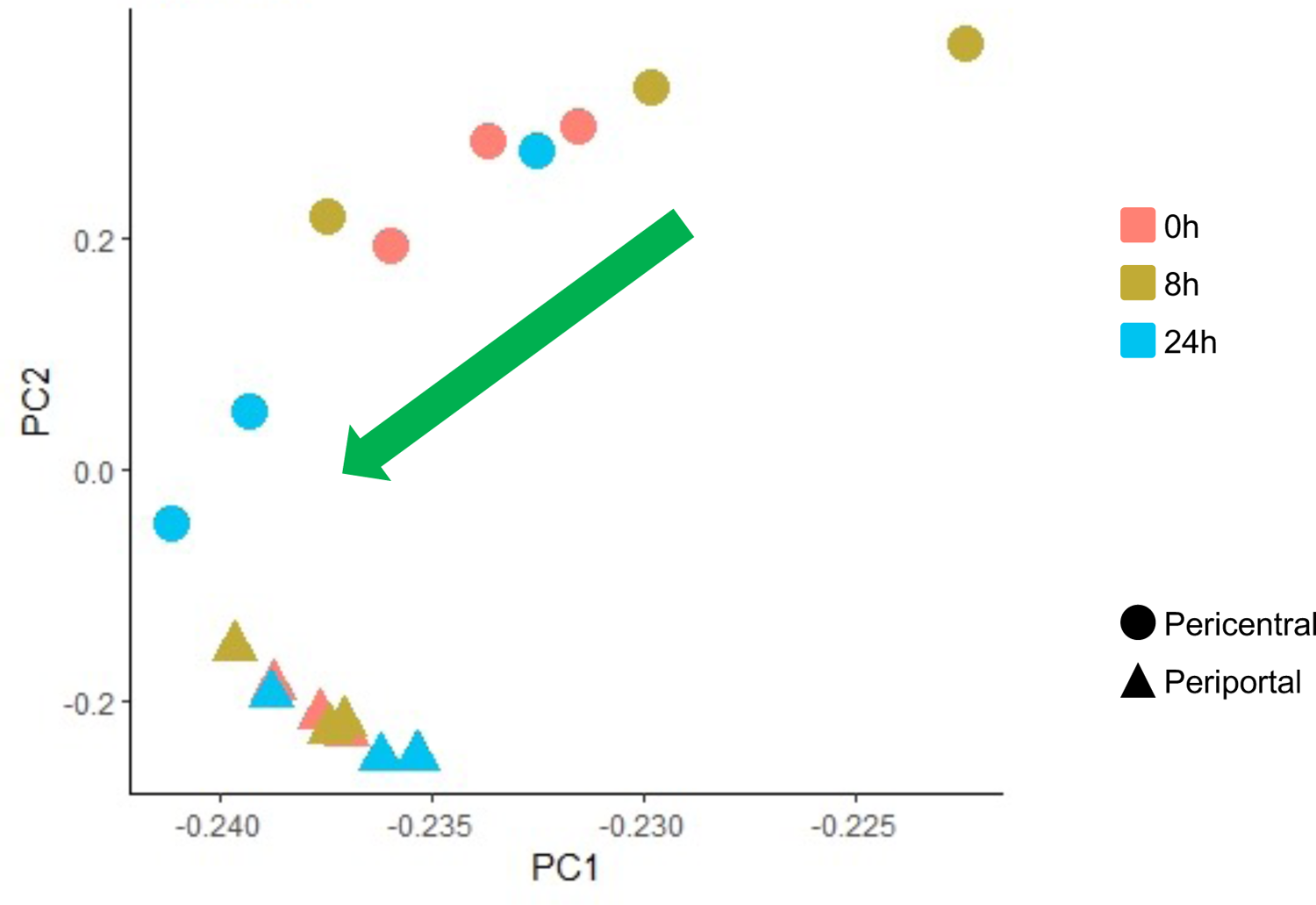
PCA of RNA-seq. Each dot represents each time point/condition based on coloring and shape of dot (0h=red, 8h=yellow, 24h=blue; circle=pericentral, triangle=periportal). Green arrow shows how fasting shift the pericentral hepatocytes.

